# Oxytocin Activity in the Paraventricular and Supramammillary Nuclei of the Hypothalamus is Essential for Social Recognition Memory in Rats

**DOI:** 10.1101/2022.05.23.493099

**Authors:** Keerthi Thirtamara Rajamani, Marie Barbier, Arthur Lefevre, Kristi Niblo, Nicholas Cordero, Shai Netser, Valery Grinevich, Shlomo Wagner, Hala Harony-Nicolas

**Affiliations:** Department of Psychiatry, New York, NY, USA; Seaver Autism Center for Research and Treatment at the Icahn School of Medicine at Mount Sinai, New York, NY, USA; Department of Neuropeptide Research in Psychiatry, Central Institute of Mental Health, Medical Faculty Mannheim, University of Heidelberg, Mannheim, Germany; Sagol Department of Neurobiology, University of Haifa, Israel; Department of Neuroscience, New York, NY, USA; Friedman Brain Institute, New York, NY, USA; Mindich Child Health and Development Institute at the Icahn School of Medicine at Mount Sinai, New York, NY, USA

## Abstract

Oxytocin plays an important role in modulating social recognition memory. However, the direct implication of oxytocin neurons of the paraventricular nucleus of the hypothalamus (PVH) and their downstream hypothalamic targets in regulating short- and long-term forms of social recognition memory has not been fully investigated. In this study, we employed a chemogenetic approach to target the activity of PVH oxytocin neurons in male rats and found that specific silencing of this neuronal population led to an impairment in short- and long-term social recognition memory. We combined viral mediated fluorescent labeling of oxytocin neurons with immunohistochemical techniques and identified the supramammillary nucleus (SuM) of the hypothalamus as a target of PVH oxytocinergic axonal projections in rats. We used multiplex fluorescence in-situ hybridization to label oxytocin receptors in the SuM and determined that they are predominantly expressed in glutamatergic neurons, including those that project to the CA2 region of the hippocampus. Finally, we used a highly selective oxytocin receptor antagonist in the SuM to examine the involvement of oxytocin signaling in modulating short- and long-term social recognition memory and found that it is necessary for the formation of both. This study discovered a previously undescribed role for the SuM in regulating social recognition memory via oxytocin signaling and reinforced the specific role of PVH oxytocin neurons in regulating this form of memory.

## INTRODUCTION

Social recognition memory (SRM) is a fundamental component of social behavior, which sub-serves everyday life interactions and is conserved across several species, including rodents^1–3^. A key feature of SRM is the ability of a species to acquire, remember and recall identities of conspecifics. This facet of cognition serves to maintain and facilitate social organizational structures among conspecifics^4^. Deficits in social recognition are a core feature of several psychiatric disorders, including autism spectrum disorder (ASD) and schizophrenia^5–7^. As such, gaining insights into the brain circuitry that drives SRM is key to understanding the neurological basis of these disorders.

Several studies have identified the neuropeptide oxytocin (OXT) as a major modulator of SRM in rodents^8–10^. OXT is produced in the paraventricular, the supraoptic, and the accessory nuclei of the hypothalamus (PVH, SON, and AN, respectively). All these nuclei project predominantly to the posterior pituitary gland, where OXT is released into the bloodstream to modulate peripheral activities such as milk ejection during breast feeding and uterus contraction during parturition^11^. In addition, PVH-OXT neurons project to a wide range of cortical and limbic structures including the hippocampus, medial amygdala, and the lateral septum, all of which are characterized by high levels of oxytocin receptor (OXTR) expression^12^ and are part of the “social recognition memory circuit”^13–16^. Collectively, this knowledge leads to the assumption that neuronal activity of PVH-OXT neurons is critical for SRM, which to date has not been directly examined.

OXT fibers and OXTR expression were previously identified in the supramammillary nucleus of the hypothalamus (SuM)^17–20^, a caudal hypothalamic nucleus that is positioned superior to the mammillary body^21^. The SuM is known to regulate hippocampal theta oscillations^22,23^ and to be involved in arousal^24^, REM sleep^25^, lactation^17^, reinforcement learning and motivation^26–30^. Recent work in mice has also demonstrated a role of the SuM in processing social novelty information^31^. Specifically, the SuM plays a role in relaying contextual and socially salient information to the hippocampus through anatomically segregated populations of projection neurons; SuM to hippocampal CA2 (SuM⍰CA2) projecting neurons process socially salient information, whereas SuM to dentate gyrus (SuM⍰DG) projecting neurons carry context specific information. The abundance of OXT fibers and receptors in the SuM, along with its role in social novelty processing, suggest that OXT signaling in the SuM is likely to regulate SRM.

In the present study, we used targeted chemogenetic inhibition to examine the specific role of PVH-OXT neurons in SRM in male rats. We then conducted viral tracing and immunohistochemical staining to identify OXT projections fibers in the SuM and complemented those with in situ hybridization to examine the presence and distribution of OXTR within SuM neurons. Additionally, we combined retrograde labeling to identify CA2 projecting SuM neurons and further examined the distribution of OXTRs across this population of neurons. Finally, we used a highly specific OXTR antagonist to examine if OXTR signaling in the SuM is crucial for SRM.

## RESULTS

### Chemogenetic silencing of PVH-OXT neurons impairs short and long-SRM

To test the direct role of PVH-OXT neurons in SRM, we examined if inhibition of PVH-OXT neuronal activity affects short and/or long-term social recognition memory following the experimental design in **Fig. 1a**. For this purpose, we used chemogenetic inhibitory DREADDs designed and validated to specifically express in OXT neurons and demonstrated to reduce mean frequency and amplitude of OXT neurons (AAV1/2-OXTp-hM4Dgi-mCherry)^32^. We first confirmed and validated previous findings indicating the specific expression of the AAV1/2-OXTp-hM4DGi-mCherry virus in PVH-OXT neurons^32^ (**Supplemental data Fig. 1a & 1b**). Next, we tested the impact of chemogenetic silencing of PVH-OXT neurons on SRM, using the social discrimination task^14,36^. When tested for short-term SRM (**Fig. 1b**), we found that rats injected with the inhibitory DREADDs showed a significant preference for the novel over the familiar social stimuli after saline (control) injection but failed to show a similar preference following CNO injection (**Fig. 1c & 1d**). Since CNO was administered prior to the initial interaction, we examined if this impaired the investigation time during the 1^st^ encounter (time when the test rat interacts with the social stimulus for the first time, as shown in **Fig. 1b**). We compared the total investigation time following CNO and saline injection and found no difference between the two treatment conditions (**Fig. 1e**). Typically, SD rats engage in different lengths of bouts during the social discrimination task and show specific temporal dynamics that are distinct from other outbred rats and mice^45^. Additionally, bouts that are shorter than 6 secs produce no clear separation of preference for the novel vs. familiar social stimuli, while bouts that are longer than 6 sec reflect more meaningful interactions in both mice^40^ and rats^45^. Therefore, we further analyzed the data based on bout lengths. As expected, we found that during short bouts (≤6sec) rats did not show preference to the novel stimuli, regardless of the treatment (saline or CNO) (**Supplemental data, Fig. 1c**). During long bouts (≥6sec), however, there was a significant preference for the novel over the familiar social stimuli following saline, but not CNO injection (**Supplemental data, Fig. 1d**).

**Figure 1.**
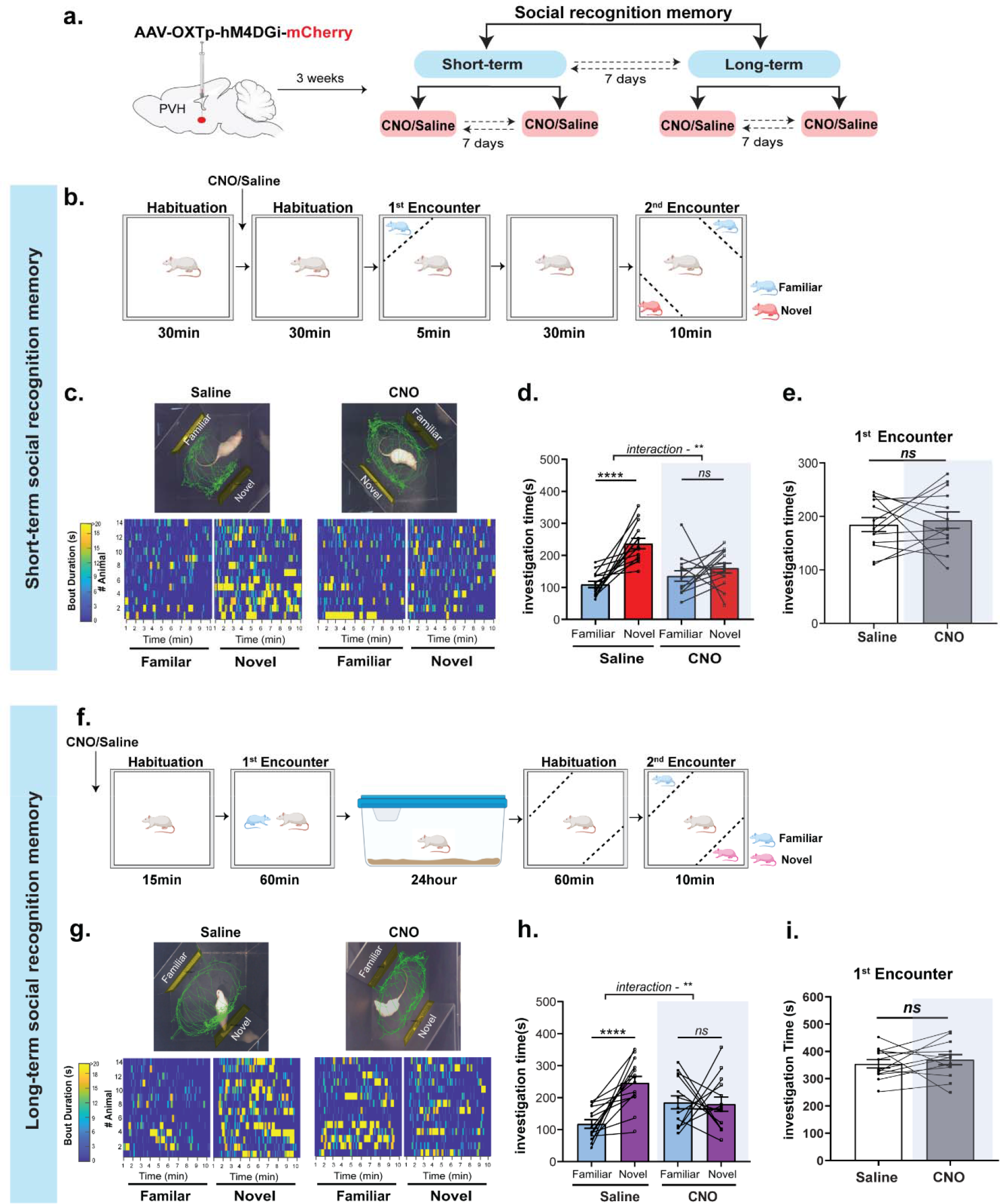
Chemogenetic silencing of PVH-OXT neurons impairs short- and long-term SRM. **a**. A schematic showing the behavioral experimental design. Saline and CNO treatments were counterbalanced between test days, and short and long-term SRM were counterbalanced between cohorts. **b**. A schematic of the experimental design for the short-term SRM. Saline or CNO was injected 30min prior to the 1^st^ encounter. **c. Top**: A representative trace (from one rat per treatment during the 2^nd^ encounter) following saline or CNO injection. **Bottom**: Heat maps representing investigation time across all rats during Novel or Familiar stimuli investigation following Saline or CNO. Each row represents one rat. **d**. Total investigation time of the Novel vs. Familiar stimuli during the 2^nd^ encounter. Saline injected rats showed a clear preference for Novel over Familiar stimuli, whereas same rats injected with CNO showed no preference for either stimuli (two-way repeated measures (RM) ANOVA, social preference (Familiar vs. Novel) x treatment (Saline vs. CNO) interaction (F_1,26_=9.11, ^**^*P*=0.0056, n=14), effect of social preference (F_1,26_ = 35.07, \*\*\*\**P*<0.0001), and effect of treatment (F_1,26_ = 1.7, *P*=0.203), post-hoc, Sidak multiple comparison test, Saline (Familiar vs. Novel, \*\*\*\**P*<0.0001) and CNO (Familiar vs. Novel, *P*=0.38, *ns*). **e**. Total investigation time of social stimuli during the 1^st^ encounter. There was no significant difference in the total investigation time between Saline and CNO treatment groups during the 1^st^ encounter (two-tailed paired student’s *t-*test, t_12_=0.45, *P*=0.65, *ns*). **f**. A schematic of the experimental design for long-term SRM. Saline or CNO was injected 15min prior the 1^st^ encounter. **g**. same as (c) but for long-term SRM**. h**. Total investigation time of the Novel vs. Familiar stimuli during the 2^nd^ encounter. Saline treated animals showed a clear preference for Novel over Familiar stimuli, whereas same rats injected with CNO showed no preference for either stimuli (two-way RM ANOVA, social preference (Familiar vs. Novel) x treatment (Saline vs. CNO) interaction (F_1,26_ = 10.51,\*\**P*=0.0032, n=14), effect of social preference (F_1,26_ =12.34, \*\**P*=0.0016), and effect of treatment (F_1,26_ = 0.0005, *P*=0.98). post-hoc, Sidak multiple comparison test, Saline (Familiar vs. Novel, ^****^*P*<0.0001) and CNO (Familiar vs. Novel, *P*=0.973, *ns*). **i**. Investigation time of the social stimuli during the 1^st^ encounter. There were no significant differences between saline and CNO injection groups during the 1^st^ encounter (two-tailed paired student’s *t-*test, t_12_=0.67, *P*=0.51, *ns*)

To examine if the effects of PVH-OXT neural inhibition on social preference is consistent across the length of the social discrimination task, we also examined social preference as a function of time. We found that following saline injection, rats maintained their preference for the novel stimuli across time whereas following CNO injection, they showed no clear preference for either stimulus along time (**Supplemental data, Fig.1e & Fig. 1f**). Furthermore, to rule out any non-specific effect of CNO on short-term SRM, and confirm that the decrease in social preference is due to an effect of the inhibitory DREADD, we injected an independent group of rats with a control virus that has the same backbone as the inhibitory DREADD virus, but lacks the hM4DGi receptor (AAV1/2-OXTp-mCherry). We first confirmed its overlap with OXT neurons (**Supplemental data Fig. 2a & 2b**), and then followed the same experimental design as described in **Fig.1b**. We found that rats showed a significant preference for the novel over the familiar social stimuli following either saline or CNO injection (**Supplemental data, Fig. 2c**), thus ruling out a non-specific effect of CNO on SRM. Finally, in order to confirm that the effect of PVH-OXT neuronal inhibition is specific for SRM and not to other aspects of non-social memory, we assessed a separate cohort of rats for their object recognition memory, using the novel object recognition memory task (**Supplemental data, Fig. 2d**). We found that OXTp-hM4DGi injected rats showed a clear preference for the novel over the familiar object, following saline or CNO injection (**Supplemental data, Fig. 2e**). Furthermore, there was no difference in the total investigation time following saline or CNO injection during the 1^st^ encounter (**Supplemental data, Fig. 2f**) and the rats showed no innate preference for one object over the other, as reflected by the similar times of investigation towards the cone or the Lego (**Supplemental data, Fig. 2g**). Taken together, these results demonstrate that the activity of PVH-OXT neurons is necessary for short-term social recognition memory.

**Figure 2.**
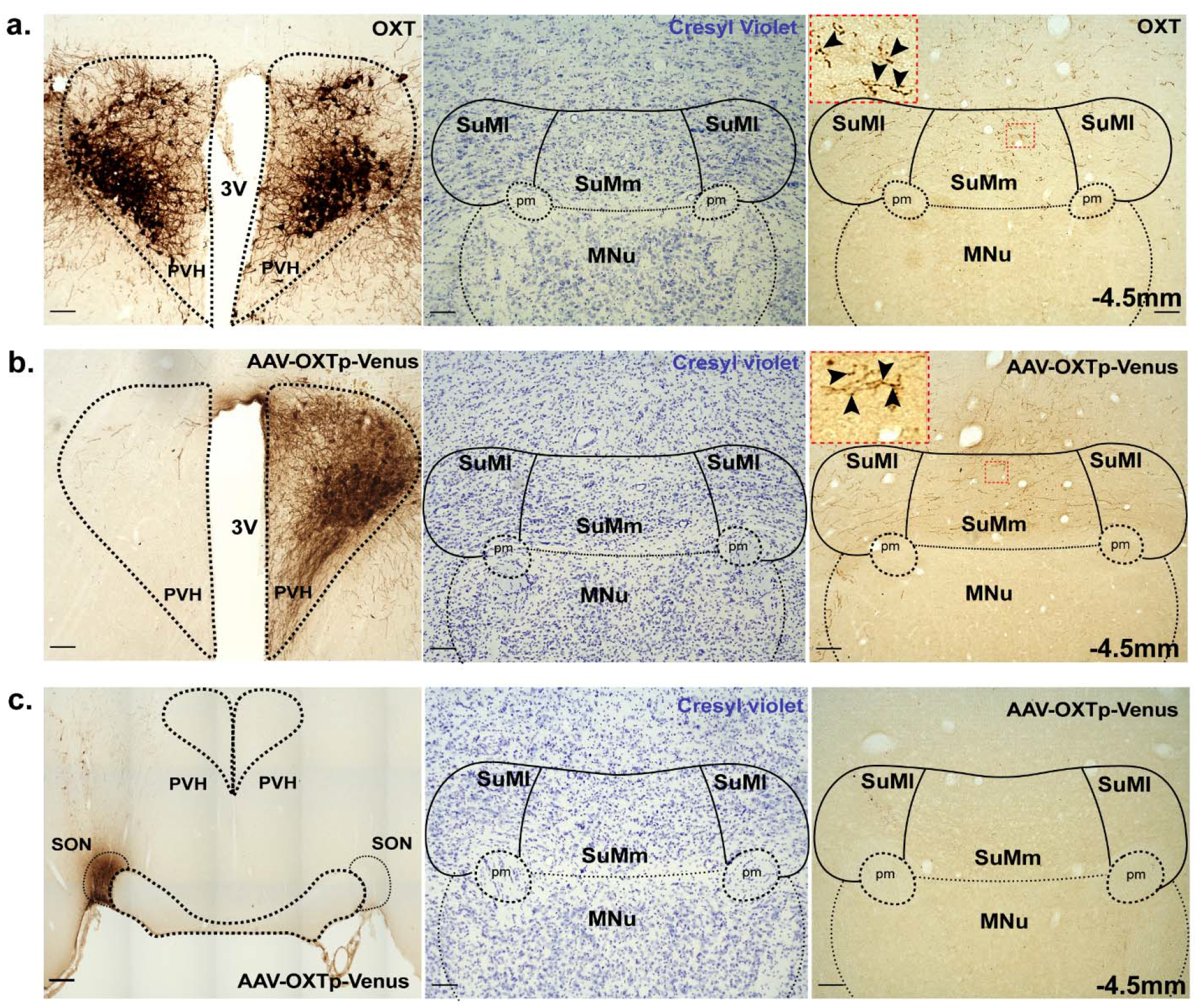
OXT fibers within the SuM originate from the PVH and not the SON. **a**. Immunohistochemical staining for OXT in the PVH and SuM**. Left**: A representative PVH section stained with specific anti OXT antibodies and developed using diaminobenzidiene (DAB) based enzymatic staining. **Middle**: A representative SuM section from the same animal as above was stained for cresyl violet to highlight anatomical structures. **Right**: An immediately adjacent SuM section to that in the middle image stained with anti-OXT antibody to highlight OXT fibers in the SuMm and the SuMl (4x). Inset shows a higher magnification (40x) image of the SuMm (highlighted by red dotted box). Black arrows point to axonal varicosities and branching axons. **b**. Immunohistochemical staining for Venus in the PVH and SuM of rats injected with the AAV1/2-OXTp-Venus in the PVH. **Left**: A representative image of the PVH injected unilaterally with the AAV1/2-OXTp-Venus. Venus was identified using anti-GFP antibody and developed enzymatically using DAB based staining. **Middle**: Cresyl violet staining of the SuM to highlight the SuMl and SUMm. **Right**: An immediately adjacent SuM section to that in the middle image shows Venus positive fibers distributed across the SuMm and SuMl (4x). Inset shows a higher magnification (40x) image of the SuMm (highlighted by red box). Black arrows point to axonal varicosities and branching axons. **c**. Immunohistochemical staining for Venus in SON and SuM from rats injected with AAV1/2-OXTp-Venus in the SON. **Left**: AAV1/2-OXTp-Venus injected into the SON identified using anti-GFP antibody. **Middle**: Cresyl violet staining of SON injected group to highlight anatomical structures in the SuM. **Right**: An immediately adjacent SuM section to that in the middle image showing absence of Venus positive fibers across the SuMm and SuMl (4x). −4.5mm denotes position of the section relative to bregma. 3V, 3^rd^ ventricle, PVH, paraventricular nucleus of the hypothalamus. SuMm, medial supramammillary nucleus, SuMl, lateral supramammillary nucleus, MNu, Mammillary nucleus, pm, principal mammillary tract. Scale bar 100um.

To determine if OXT neurons in the PVH are also necessary for long-term SRM we tested the same cohort of rats, which were tested on the short-term social discrimination task, on the long-term social discrimination task (**Fig. 1f**). We found that following saline injection, inhibitory DREADDs-injected rats showed significant preference for the novel over the familiar social stimuli. However, the same rats failed to show such preference following CNO injection (**Fig. 1g & 1h**). Furthermore, no significant differences between treatments were observed in the investigation time during the first 10 minutes of the 1^st^ encounter (**Fig. 1i**). By classifying the stimuli interaction time into short (≤6sec) and long (≥6sec) bouts, we found that rats do not display any social preference when analyzing the short bout interactions, regardless of treatment (saline or CNO) (**Supplemental data, Fig. 3a**). However, preference to the novel stimuli over the familiar social stimuli was clearly observed when analyzing long bouts following saline, but not CNO injection (Supplemental data, **Fig. 3b**). By analyzing the social preference data across time, we found that preference to the novel stimuli was sustained throughout the duration of the testing period following saline but no clear preference was observed following CNO injection (**Supplemental data, Fig. 3c & 3d**). Altogether, these findings demonstrated that PVH-OXT neurons also play a critical role in long-term SRM and are likely to be a common substrate that mediates both short and long-term SRM.

**Figure 3.**
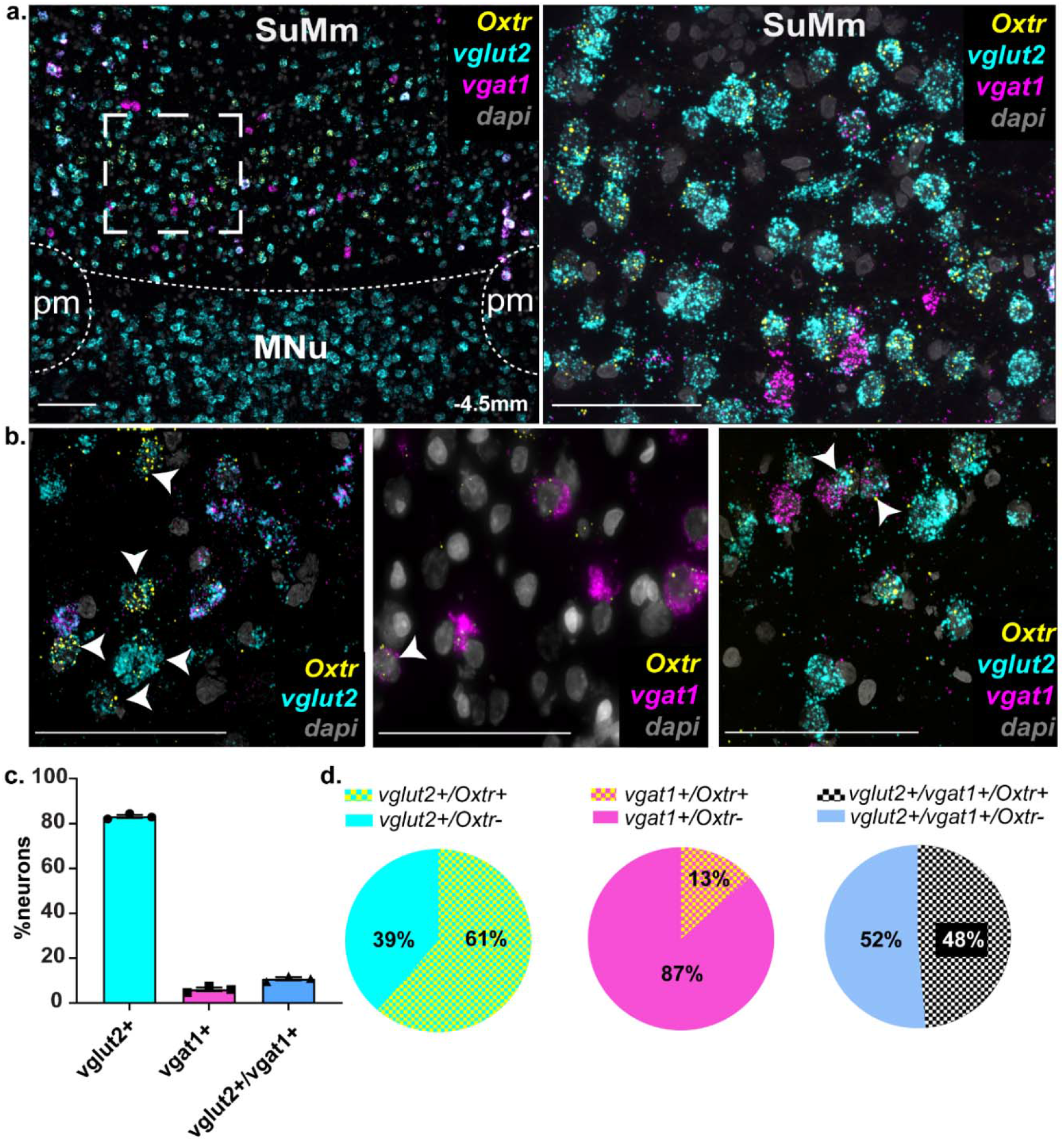
OXTRs are differentially distributed in SuM neurons. **a**. RNAscope was performed on SuM tissue using probes for *Oxtr*, *vglut2* (marker for glutamatergic neurons) and *vgat1* (marker for GABAergic neurons). **Left**: Lower magnification (10x) shows *Oxtr* along with *vglut2* and *vgat1* expression in the SuM. **Right**: Higher magnification (40x) of SuM tissue shows distribution of *Oxtr* across *vglut2^+^* and *vgat1*^+^ neurons. **b**. Higher magnification (63x) of SuM tissue showing *Oxtr* localization in *vglut2*^+^ neurons (**Left**), *vgat1^+^* neurons (**Middle**), or *vglut2^+^*:*vgat1*^+^ neurons (**Right**), highlighted by white arrows. **c**. Quantification of *vglut2*^+^, *vgat1^+^*, and *vglut2^+^:vgat1^+^* neurons in the SuM. **d**. Quantification of *Oxtr* distribution across *vglut2^+^* (**Left)**, *vgat1^+^* (**Middle**) and *vglut2^+^:vgat1* neurons (**Right**) neurons. Scale bar 100um, SuM, Supramammillary nucleus. MNu, Mammillary nucleus, pm, principal mammillary tract. Data presented as Mean±SEM.

### The supramammillary nucleus is a target for OXT innervation that originates in the PVH but not the SON

The SuM is a caudal hypothalamic nuclei that is juxtaposed immediately over the mammillary bodies^23^. It has been shown to contain OXT fibers, yet the source of these fibers has not been determined^17^. Given the recently described role of the SuM in regulating social novelty processing in mice^31^, we hypothesized that OXT action within the SuM is likely to contribute to the SuM’s role in social novelty processing and the formation of social memory. We first wanted to confirm previous findings by demonstrating the presence of OXT fibers in the SuM and determining the source of its innervation. For this purpose, we used immunohistochemistry with anti-OXT antibodies to visualize OXT fibers in the SuM and found that they are broadly distributed across the rostro-caudal parts of the SuM with fibers identified in both the medial (SuMm) and lateral part of the SuM (SuMl) (**Fig. 2a and Supplemental data, Fig. 4**). To determine the origin of these OXT fibers, we unilaterally injected, into the PVH or SON, an anterograde virus, which expresses Venus fluorescent protein specifically in OXT neurons (AAV1/2-OXTp-Venus), thus allowing specific labelling of OXT neurons and their projections^33^. We found that the PVH is a major source for OXT fibers in both the SuMm and SuMl (**Fig. 2b and Supplemental data, Fig. 4**) and observed punctate labeling as well as branched axons indicative of local innervation (**Fig. 2a & 2b**, Right, insets). We further confirmed the presence of OXT axonal terminals in the SuM by injecting an anterograde AAV-OXTp-synaptophysin-GFP virus^33^, which expresses GFP fused to synaptophysin, a presynaptic terminal marker in OXT neurons (**Supplemental data, Fig. 5a**). Calretinin and parvalbumin have been used to distinguish the SuM, which is calretinin positive but parvalbumin negative, from the mammillary body, which is located caudal to the SuM and is parvalbumin positive but calretinin negative^23^. We, therefore, used these markers in combination with anti-OXT antibodies (Supplemental Data, **Fig. 5b**) or AAV1/2-OXTp-Venus (Supplemental data. **Fig. 5c**) to confirm the high abundance of OXT fibers in the SuM, but not the mammillary body. Importantly, we did not detect any OXT fibers in the SuM when the virus (AAV1/2-OXTp-Venus) was injected into the SON (**Fig. 2c**), suggesting that the SON OXT neurons do not project to the SuM. All together, these experiments demonstrate that PVH-OXT neurons are a major source for OXT projection fibers in the SuM.

**Figure 4.**
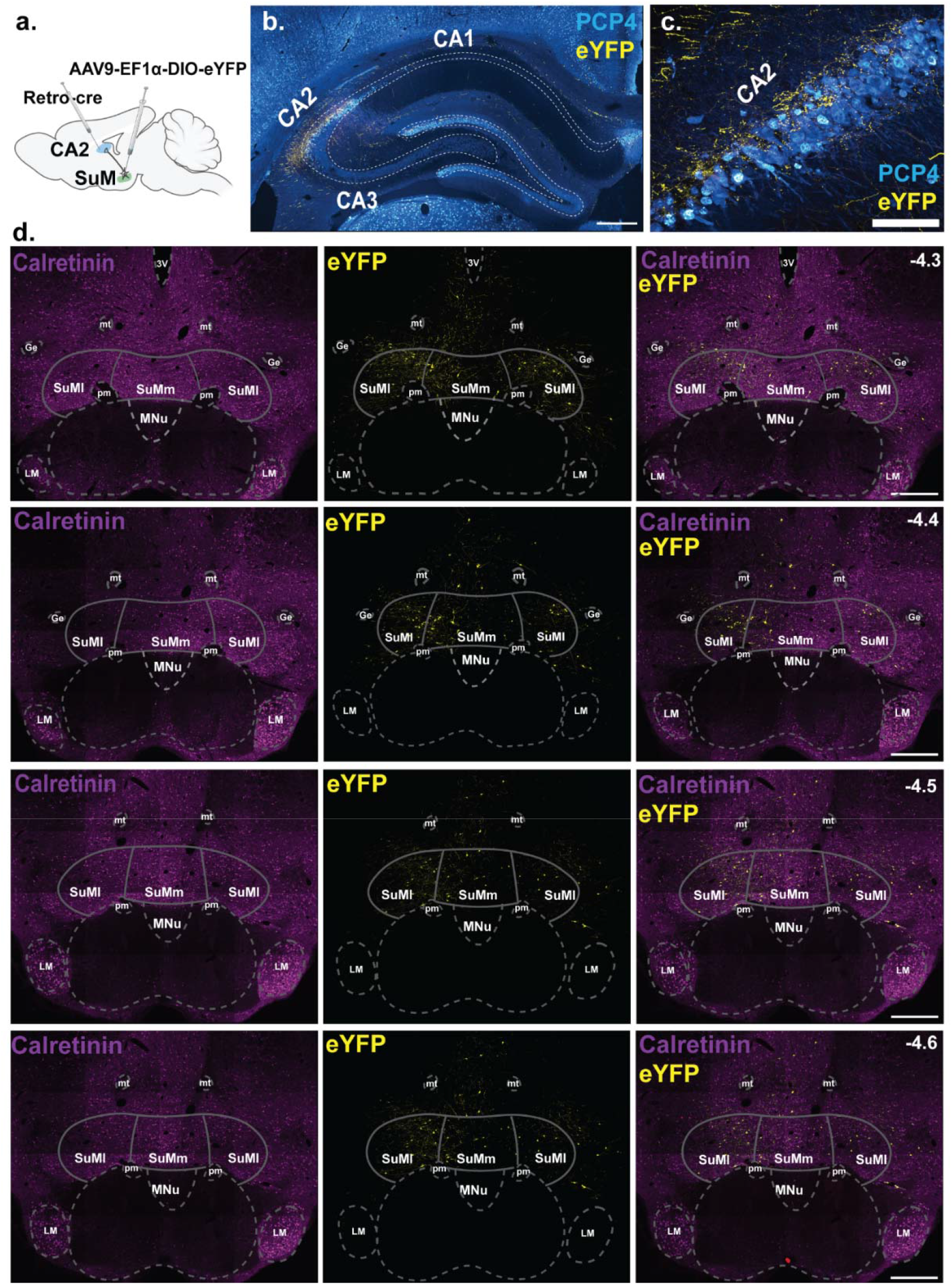
SuM neurons project to the hippocampal CA2. **a**. A schematic of the viral strategy to target and identify SuM⍰ CA2 projecting neurons. A retrograde virus expressing a cre-recombinase (AAV-Ef1α-Rg-Cre) was injected into the hippocampal CA2 and a cre-dependent anterograde virus (AAV9-EF1□-DIO-eYFP) was injected into the SuM. **b**. A coronal section show SuM axonal terminals (labelled with eYFP) in the CA2 neurons (identified by anti-PCP4 staining). **c**. A higher magnification of the hippocampal CA2. **d**. Four representative images spanning the SuM show SuM⍰ CA2 projecting neurons across the rostro-causal and dorso-ventral axis of the SuM. Calretinin was used to highlight the boundaries between the SuM and the mammillary nucleus. −4.3 to −4.6mm denotes position of the section relative to bregma. Ge, Nucleus of Gemini, mt, mamillothalamic tract, pm, principal mammillary tract, SuMl, supramammillary nucleus lateral, SuMm, supramammillary nucleus medial. MNu, Mammillary nucleus. Scale bar: 5b and 5d, 500um; 5c, 100um.

**Figure 5.**
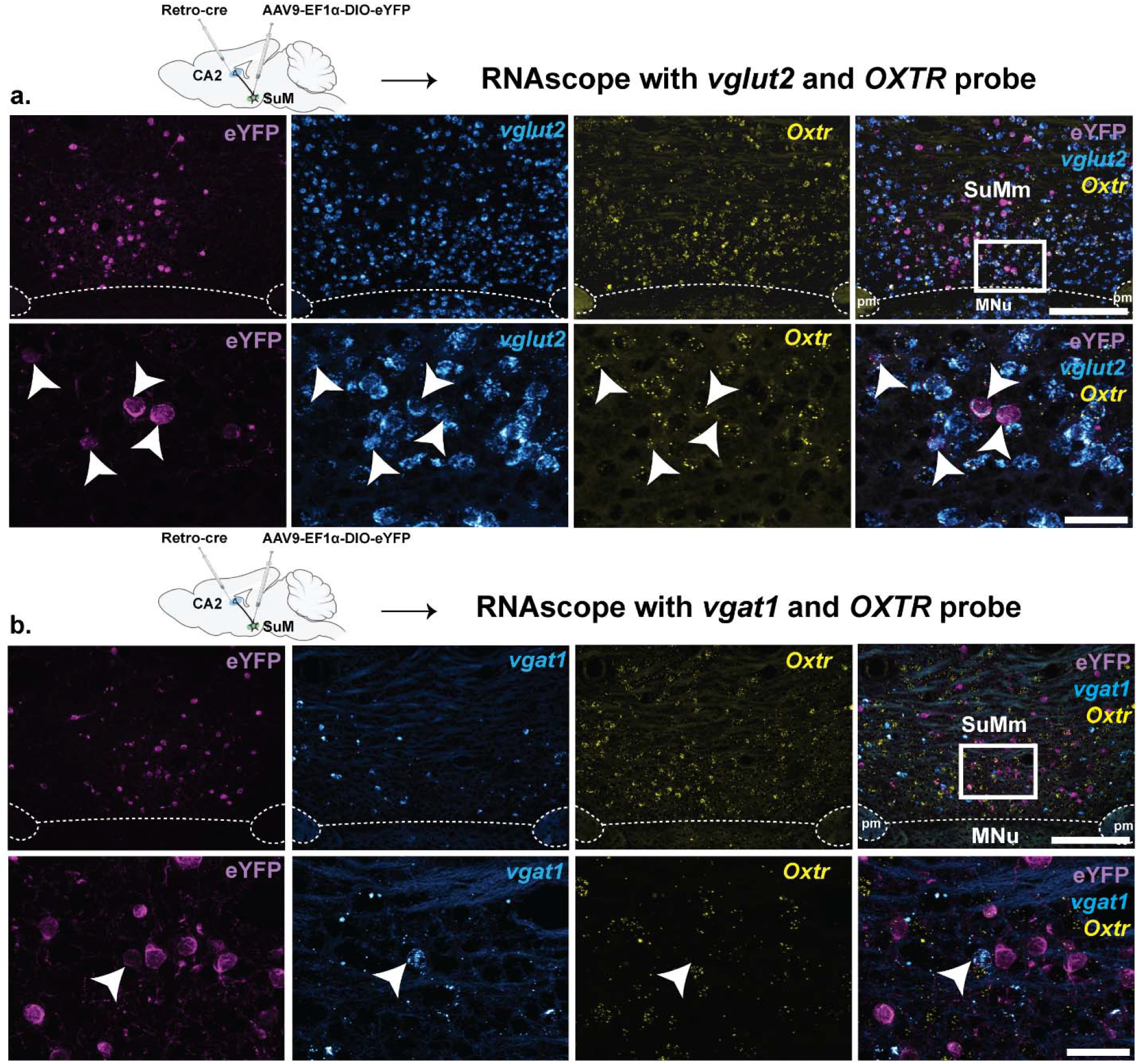
SuM^vglut2+^ ⍰ CA2 projecting neurons co-express oxytocin receptors. **a**. RNAscope using probes for *vglut2* and *Oxtr*, followed by immunohistochemistry for eYFP on tissue section from rats injected in the hippocampal CA2 with a retrograde virus expressing a cre-recombinase (AAV-Ef1α-Rg-Cre) and in the SuM with a cre-dependent anterograde virus (AAV9-EF1□-DIO-eYFP). **Top**: lower magnification of SuM showing expression of eYFP, *vglut2*, and *Oxtr* in the SuM. **Bottom**: higher magnification of the same Top images. White arrow point to the overlap between SuM⍰ CA2 projecting neurons (labeled with eYFP), *vglut2* and *Oxtr*. **b**. same as in **a**, but using the *vgat1* instead of the *vglut2* probe. White arrows point to the lack of expression of *Oxtr* in eYFP and *vgat1* positive neurons. Scale bar: 6a and 6b - Top panels, 250um. 6a, and 6b - Bottom panels, 50um.

### OXT receptors are expressed by specific population of SuM neurons

After establishing that PVH-OXT are a major source of OXT innervation in the SuM, we set to determine if SuM neurons express OXTRs. Although previous studies have identified that the SuM contains OXTR using autoradiographic techniques in rats^19,20^ or transgenic mice^18^, those studies were limited by their inability to determine the cellular distribution of the receptors. The SuM is comprised mostly of glutamatergic neurons and to a lesser extent of GABAergic, dopaminergic^46^, substance P^47^, and CCK positive neurons^48^. Additionally, the SuM is one of the few brain regions in the rat where some neurons co-express both glutamate and GABA^43^. In order to determine the cellular distribution of OXTR within the SuM, we employed RNAscope, an *in situ* RNA hybridization (ISHr) technology, to identify OXTR transcripts (*Oxtr)* and examine their overlap with glutamatergic and GABAergic neurons. We used the vesicular glutamate transporter (*vglut2)*, as a marker for excitatory neurons, and the vesicular GABA transporter (*vgat1)* as a marker for inhibitory neurons **(Supplemental data Fig. 6a)**. We first examined the proportion of SuM neurons that are GABAergic, glutamatergic, or both and then determined if *Oxtr* differentially segregate across these neural populations. We found that neurons within the SuM are primarily positive for *vglut2* (83±0.7%, # of *vglut2*^+^ neurons, 344±2.3, total number of neurons, 413.6±2.72; n=3 rats, 1 section/rat), with a small population being positive for both *vglut2* and *vgat1* (10.7±0.7%, # of *vglut2^+^:vgat1^+^* neurons, 44.6±3.17), and an even smaller population of neurons that were positive only for *vgat1* (6±0.8%, # of *vgat1^+^* neurons, 25±3.4) (**Fig. 3a-c**). In order to determine how *Oxtr* segregates into these populations, we quantified the number of neurons that express *Oxtr* within each population. We found that nearly 60% of *vglut2*^+^ neurons are also *Oxtr*^+^ (# of *vglut2*^+^/*oxtr*^+^, 211±25.1), whereas 48% of *vglut2*^+^:*vgat1*^+^ neurons are *Oxtr*^+^ (# of *vglut2^+^:vgat1^+^/Oxtr^+^* neurons, 21.6±1.2) and only 13% of *vgat1*^+^ are *Oxtr*^+^ # of *vgat1*^+^/*Oxtr*^+^ neurons, 3.3±0.8 (**Fig. 3d**). These results indicate that OXTRs in the SuM are predominantly expressed in glutamatergic neurons and neurons that co-express glutamate, and GABA and in a very minor population of GABA neurons.

### SuM^vglut2+^ neurons that project to the hippocampal CA2 area co-express OXTR

SuM neurons have been shown to project to the hippocampus, specifically the hippocampal CA2 in mice^31,49^. Here, we set to confirm if SuM⍰CA2 projections also exist in rats and to determine if OXTRs are specifically expressed in these projecting neurons. To this end, we injected a retrograde virus (AAV-Ef1α-Rg-Cre) into the hippocampal CA2 and a Cre-dependent reporter virus (AAV-Ef1α-DIO-eYFP) into the SuM (**Fig. 4a)**. The AAV-Ef1a-DIO-eYFP virus travels anterogradely by virtue of its packaging capsid (AAV9) (**Fig. 4a**), thus the combination of the two viruses allowed us not only to identify CA2 projecting neurons in the SuM, but also to visualize axonal terminals of these neurons in the CA2 (**Fig. 4b-c)**. We identified SuM⍰CA2 projections neurons across the rostro-caudal and dorso-ventral axis of spread (Bregma, −4.3 to −4.6mm) and in both the medial and the lateral boundary of the medial and the lateral aspect of the SuM (**Fig. 4d**). In order to determine if OXT receptors are expressed by this subset of neurons, we performed ISHr for *Oxtr* and *vglut2* or *vgat1 and* stained for GFP using immunohistochemistry and found O*xtr* transcripts to be co-localized in SuM^*vglut2*+/GFP+^⍰CA2 projecting neurons (**Fig. 5a**) but not on SuM^*vgat1*+/GFP+^ neurons (**Fig. 5b**). Taken together, these results indicate that OXTRs are localized on excitatory SuM neurons that project to the hippocampal CA2 and suggest that activity of this neuronal population could be modulated by OXTR signaling.

### Blocking OXTR in the SuM affects both short and long social recognition memory

To follow up on our findings, which indicated that the SuM is heavily innervated by PVH-OXT fibers and exhibits high expression of OXTR, we set to test if OXT downstream signaling within the SuM is necessary for SRM. To this end, we implanted a cannula to target the SuM (**Supplemental data Fig. 7**) and infused a selective OXTR antagonist (0.3ul, 75ng total)^34,35^, 10 min before testing the rats on the short or long-term social discrimination tasks, using a cross-over design (**Fig. 6a**). We found that on the short-term SRM task (**Fig. 6b**), following saline infusion, subject rats showed a clear preference for the novel over the familiar social stimuli, whereas infusion of OXTR antagonist led to a lack of preference for either stimuli (**Fig. 6c and 6d)**. Furthermore, we found no significant differences in investigation time between the saline and OXTR antagonist-infused rats during the 1^st^ encounter (**Fig. 6e)**. As before, when the short bouts were assessed, neither saline nor OXTR antagonist injected rats showed any preference for the novel stimuli (**Supplemental data, Fig. 8a**). However, when bouts that were longer than 6 secs where assessed, saline injected rats showed a clear and significant preference for the novel over the familiar stimuli, with no such preference observed following OXTR antagonist infusion (**Supplemental data, Fig. 8b**). The preference for the novel stimuli over the familiar stimuli in the saline infused rats and the lack of clear preference in the OXTR infused rats were sustained across time (**Supplemental data**, **Fig. 8c & 8d**). Overall, these results suggest that OXT signaling in the SuM is necessary for modulating short-term SRM.

**Figure 6.**
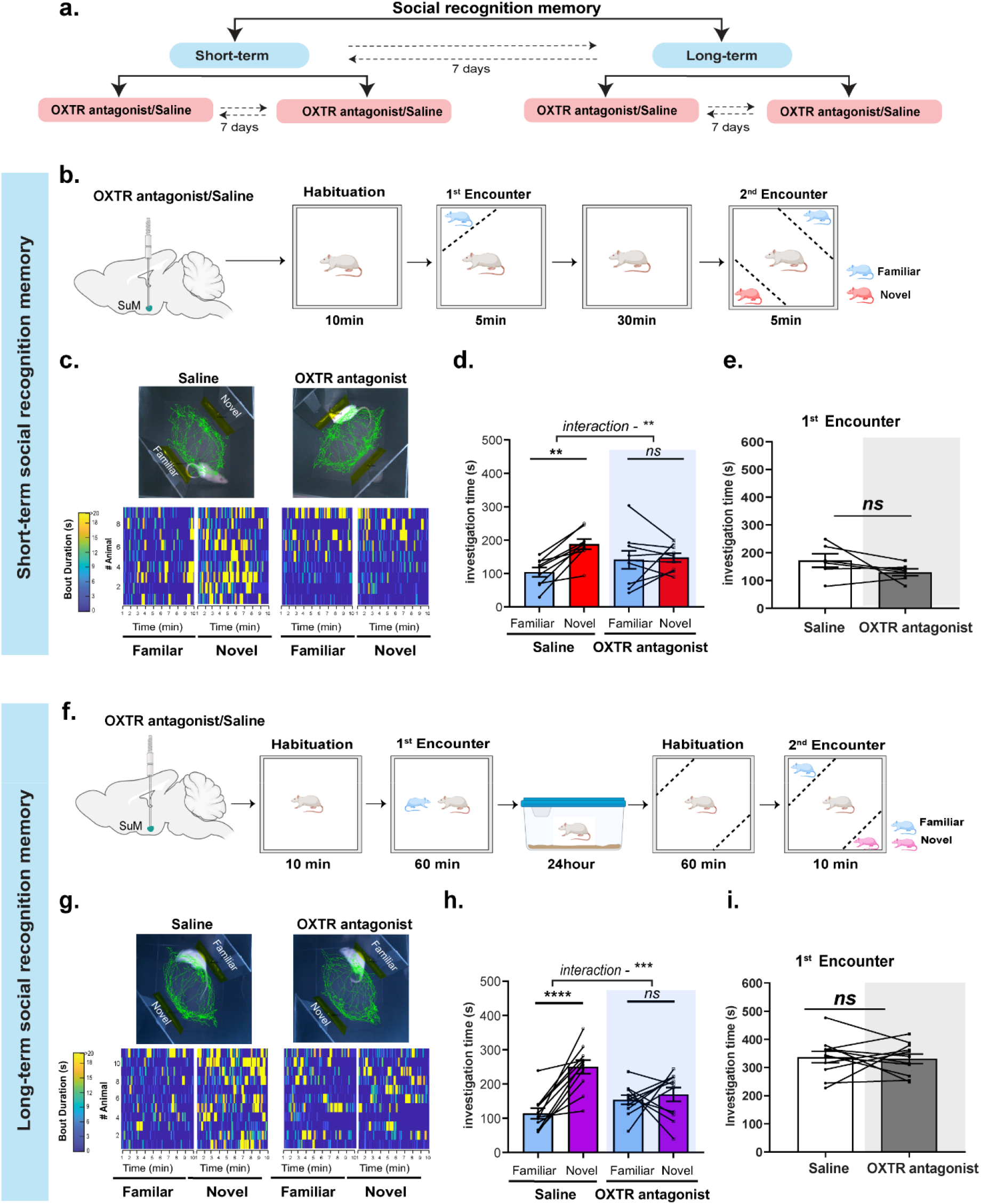
OXTR antagonism in the SuM affects short and Long-term SRM. **a**. A schematic showing the behavioral experimental design. Saline and OXTR antagonist treatment were counterbalanced between test days, and short and long-term SRM was counterbalanced between cohorts. **B**. A schematic of the behavioral paradigm for short-term SRM. Saline or OXTR antagonist was infused 10min prior to the 1^st^ encounter. **c. Top**: A representative trace from one animal per treatment during the 2^nd^ encounter following Saline or OXTR antagonist infusion. **Bottom**: Heat maps representing investigation time of all rats during novel or familiar investigation following saline or OXTR antagonist infusion. **D**. Total investigation time of the novel vs. familiar stimuli during the 2^nd^ encounter. Saline infused rats showed a clear preference for Novel over Familiar stimuli, whereas the same rats showed no preference for Novel or Familiar stimuli after OXTR antagonist infusion (two-way RM ANOVA, social preference (Familiar vs. Novel) x treatment (Saline vs. OXTR antagonist) interaction (F_1, 16_ = 9.25, *\*\*P*=0.007, n=9), effect of social preference (F_1, 16_ = 4.02, *P*=0.06), and effect of treatment (F_1, 16_ = 0.01, *P*=0.89). Post-hoc, Sidak multiple comparison test, Saline (Familiar vs. Novel) ^**^*P*=0.005, OXTR antagonist (Familiar vs. Novel, *P*=0.96, *ns*). **e**. Investigation time of the social stimuli during the 1^st^ encounter. There was no significant difference between saline and OXTR antagonist treatment during the 1^st^ encounter (two-tailed student’s *t* test, t_5_=1.513, *P*=0.19, *ns*). **f**. A schematic of the behavioral paradigm for long-term SRM. Saline or OXTR antagonist was infused 10min prior to the 1^st^ encounter. **g. Top**: same as (c) but for long-term SRM **h**. Total investigation time of the novel vs. familiar stimuli during the 2^nd^ encounter. Saline infused rats showed a clear preference for novel over familiar stimuli, whereas the same rats showed no preference for the either stimuli after OXTR antagonist infusion (Two-way RM ANOVA, social preference x treatment (Saline v OXTR antagonist) interaction, F_1,20_ = 15.32, \*\*\**P*=0.0009, n=11), effect of social preference (F_1,20_ = 15.27, \*\*\**P*=0.0009), and effect of treatment (F_1,20_ = 1.723, *P*=0.2). Post-hoc, Sidak multiple comparison test, Saline (Familiar vs. Novel, ^****^*P*<0.0001) and OXTR antagonist (Familiar vs. Novel, *P*=0.78, *ns*). **i**. There was no significant difference between saline and OXTR antagonist infused group during the 1^st^ encounter on the long-term SRM (two-tailed student’s *t* test, t_10_=0.24, *P*=0.80, *ns*).

Similarly, we examined the impact of the OXTR antagonist on long-term SRM (**Fig. 6f**). We found that compared to saline infusion, where rats showed a clear preference for the novel over the familiar stimuli, rats infused with OXTR antagonist in the SuM showed a robust impairment in long-term SRM (**Fig. 6g & 6h**). Finally, there was no significant difference in the investigation time during the first 10 minutes of the 1^st^ encounter between the saline and OXTR infused groups (**Fig. 6i**). As expected, short interaction bouts showed no significant differences in the preference for the novel over the familiar social stimuli, regardless of whether rats were infused with saline or OXTR antagonist (**Supplemental data, Fig. 8f**). When long interaction bouts were assessed, saline infused rats showed a clear preference for the novel over the familiar stimuli, whereas OXTR antagonist infused rats failed to show a preference for either stimuli (**Supplemental data, Fig. 8g**). The preference for the novel stimuli over the familiar stimuli in the saline infused rats and the lack of clear preference in the OXTR infused rats were both sustained over time (**Supplemental data, Fig. 8h and 8i**).

## DISCUSSION

Social recognition memory is a key component of social behavior that is essential for distinguishing between familiar and novel conspecifics^50,3^ and is regulated by a defined brain circuit^2,3^ This circuit engages several neural substrates including the lateral and medial septum, prefrontal cortex, medial amygdala and hippocampus^14,51–53^. Information processing within neural circuits is not hard-wired but rather adaptive to the surrounding environment, in part due to the activity of neuromodulators such as OXT^54–56^.

In this study, we showed that acute silencing of OXT neurons within the PVH impaired the ability of rats to discriminate between novel and familiar social stimuli, indicating that activity of PVH-OXT neurons is critical for mediating SRM in rats. These findings align with the previously established role of OXT in short and long-term SRM^2,8,10,15,36,51,57^ yet attributes, for the first time, a specific role for PVH-OXT in both forms of memory. In contrast to a recent study in male mice showing that chemogenetic silencing of PVH-OXT neurons decreases social investigation of a first-time presented (novel) social stimulus^58^, our study further demonstrated that acute chemogenetic silencing of PVH-OXT neural activity in rats had no impact on social investigation of a novel social stimulus. Our findings align with discoveries from previous studies, which assessed social investigation and SRM in male mice and prairie voles that lacked the OXT or OXTR encoding gene^9,10,59^. Male OXT-KO or OXTR-KO mice and their WT and littermates spent similar amount of time investigating a first-time presented stimulus in a free interaction setup, indicating an intact approach and interest to engage in social interaction. However, when exposed repeatedly to the same stimulus, OXT-KO and the OXTR-KO mice failed to form SRM^9,10^. In prairie voles, OXTR-KO males showed no deficits in investigating a novel stimulus on the three-chamber test, but showed deficits in SRM when presented with a novel and familiar stimulus^59^. These studies, together with our findings, suggest that OXT may not be essential for the act of social interaction per se, but rather necessary for the formation of SRM, which involves processing of sensory information, encoding the salience of social stimuli, and forming social memory. Notably, a recent study in female rats showed that specific silencing of the parvocellular population of PVH-OXT neurons led to a decrease in the investigation time of a social stimulus, but only during a free and not a contained social interaction^60^. Further investigation of these findings led the authors to conclude that social touch promotes social communication in female rats through the activity of the parvocellular PVH-OXT neurons. These findings raise an important question regarding the role of OXT and social touch, as well as other sensory modalities in salience encoding during social interaction and the formation of SRM, which ought to be addressed in future studies. The direct implication of PVH-OXT neurons in modulating social memory is of significance as rodent models with mutations in high-risk genes for autism spectrum disorder (ASD), have shown changes in the overall number of PVH-OXT neurons and/or reduced OXT levels, thus suggesting that modified OXT activity could underlie some of the social behavioral phenotype reported in these models^58,61,62^. Our own work in a rat model that harbors a mutation in a high-risk gene for ASD, *Shank3*, identified long-but not short-term SRM deficits that could be ameliorated with exogenous administration of OXT^36^, thus reinforcing the need to dissect the role of OXT in modulating SRM.

The SuM is involved in several functions including arousal^24^, REM sleep^25^, lactation^17^, locomotion^63^, reinforcement learning and motivation^26–30^. SuM activity is also important for synchronizing the frequency of theta activity in the hippocampus, which modulates spike-time coordination during spatial navigation to regulate spatial memory^23,63–65^. Recently, Chen et al, demonstrated that the SuM is involved in processing social novelty information^31^ via a specific pathway that connects the SuM with the hippocampal CA2 (SuM⍰ CA2 pathway). They also showed that activation of the SuM⍰ CA2 pathway regulates the excitation vs. inhibition (E/I) ratio within the CA2, which in turn could play a role in tuning the response of CA2 neurons to novel social stimuli. Although OXT fibers have been previously reported in the SuM^17^, it was unclear if these fibers originated from the PVH or the SON. Our findings demonstrate that PVH-OXT neurons are a source for OXT fibers in the SuM. Furthermore, the SuM has been shown to express OXTRs in both mice^18^ and rats^19,20^, however, to our knowledge, our study is the first to show that OXTRs is localized on glutamate, GABA and glutamate/GABA co-expressing SuM neurons and that OXT receptors are expressed on glutamatergic neurons that project to the hippocampal CA2. These findings are significant considering the role of the SuM in regulating social novelty processing via the SuM⍰CA2 pathway^31^, and together with our findings, they suggested that social memory is likely to be modulated by OXT signaling in the SuM. Indeed, we found that blocking OXT signaling within the SuM impaired both short- and long-term social memory, thus providing evidence for OXT’s role within the SuM in social memory.

Based on these findings and the accumulated knowledge in the field, we propose a working model where the PVH, the SuM, and the hippocampal CA2 work together to modulate SRM. Specifically, we propose that the PVH-OXT⍰ SuM pathway acts to amplify the salience of the social stimulus via OXTR signaling within the SuM, while the SuM⍰CA2 pathway routes the social information to the CA2 to facilitate social memory (**Fig. 7**). Future studies, designed to manipulate PVH-OXT⍰ SuM or the SuM^vglut2+/OXTR+^⍰ CA2 pathway at different time points during the SRM task, will narrow down the specific contribution of each of these pathways to the acquisition, consolidation, and/or recall phase of SRM. Furthermore, studies, using in vivo and in vitro recordings in the SuM following manipulation of PVH-OXT neuronal activity or OXTR agonist/antagonist administration will shed a light onto the molecular and cellular mechanisms by which OXT exerts its effect on SuM neurons and indirectly on CA2 neurons to regulate SRM. To the best of our knowledge, Cumbers and colleagues were the only to examine the effect of OXT on SuM neuronal activity^17^. They found that OXT infusion into the SuM increased the spiking rate of a subpopulation of SuM neurons and facilitated suckling-evoked SuM neural responses in lactating rats. Building on these findings, future in vitro studies should examine the effect of OXT not only on spontaneous firing, but also intrinsic excitability, as well as inhibitory and excitability synaptic transmission in SuM neurons, which will enhance our understanding of the OXT action in the SuM. *In vivo* studies, on the other hand, could examine the role of OXT in regulating SuM and CA2 neuronal activity during SRM and test the causality between OXT signaling, SuM and CA2 neuronal activity, and SRM.

**Figure 7.**
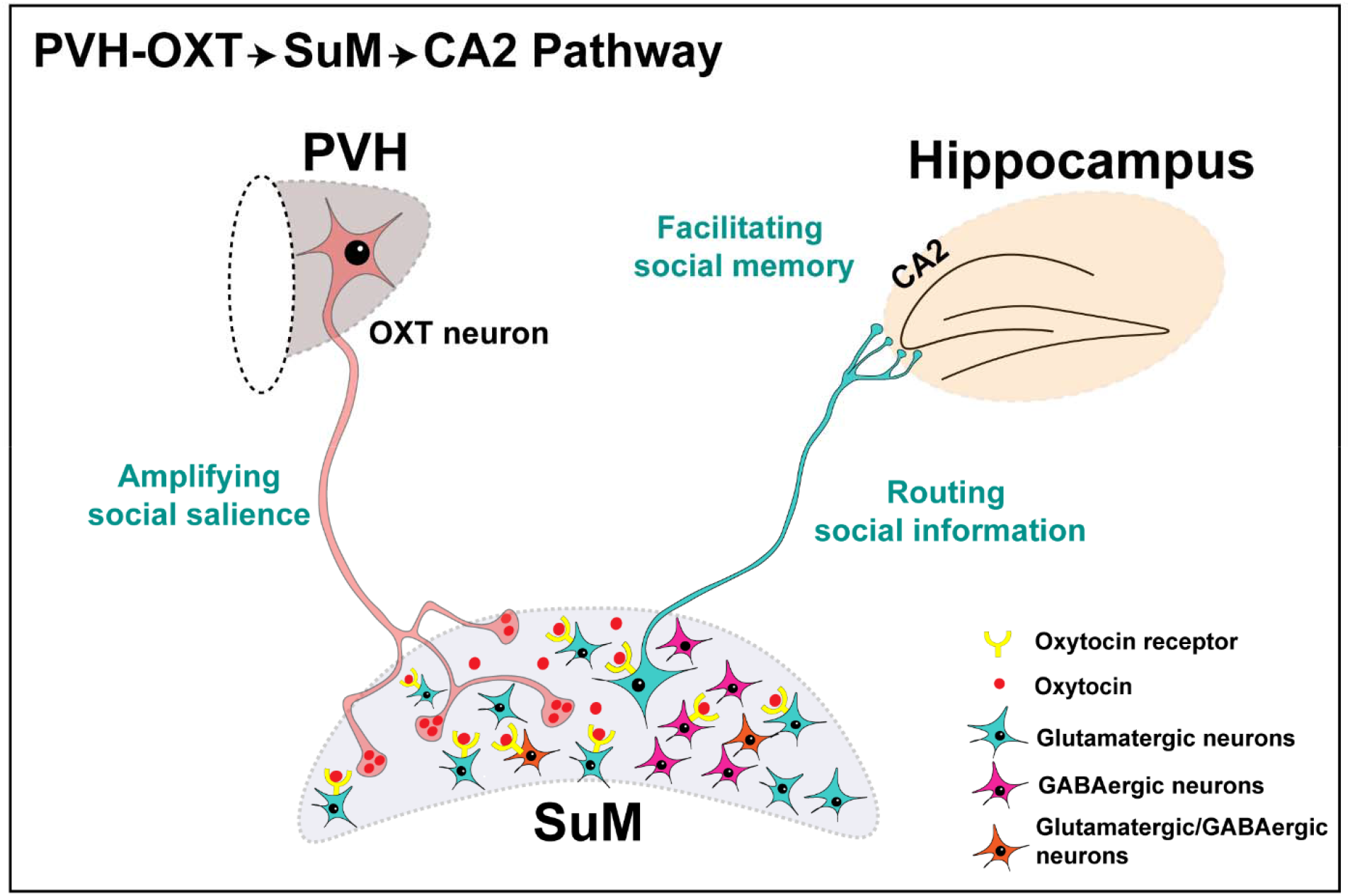
Working model of a novel neural circuit underlying social recognition memory. Our working model proposes that OXT release from PVH-OXT neuronal terminals within the SuM modulates the saliency of social signals and in turns enhances the routing of social information to the hippocampal CA2 region, where the social memory is formed. PVH, Paraventricular nucleus, OXT, oxytocin. SuM, Supramammillary nucleus.

Testing our working model will not only enhance our understanding of social brain circuits, but also has the potential to identify new targets for treatment of social behavior deficits, including deficits in SRM, which are abundant in several psychiatric disorders such as schizophrenia and ASD ^5–7^.

## METHOD DETAILS

### EXPERIMENTAL MODEL AND SUBJECT DETAILS

Male Sprague Dawley (SD) rats (Charles River, Wilmington, MA, USA) were used as test subjects for all experiments. 3-5 week old male Wistar and Wistar Hannover rats (Charles River, Wilmington, MA, USA) were used as stimuli strain for the social recognition memory experiments. All stereotaxic viral injections and cannula implantations were performed at the age of 8 weeks. Animals were housed in groups of 2 under a 12h light/dark cycle at 22 ± 2°C with food and water available *ad libitum*. All animal procedures were carried out in accordance with protocols approved by the Institutional Animal Care and Use Committee at the Icahn School of Medicine at Mount Sinai.

### Experimental Design

To take advantage of the designer receptors activated by designer drugs (DREADD) system, we used a cross-over design wherein the same rat received either 0.9% saline or clozapine N Oxide (CNO)/OXTR antagonist across the testing paradigm (**Fig. 1a, 6a**). The behavioral task is detailed here: Half of the experimental rats in the cohort received saline and the remaining received CNO/OXTR antagonist and were tested on the short-term social discrimination task to assess short-term SRM. A week later, the rats that previously received saline now received CNO/OXTR antagonist and vice versa and were tested on the short-term social discrimination task to assess short-term SRM. A week after assessing short-term SRM, the same experimental design was repeated but this time, the rats were tested on the long-term social discrimination task to assess long-term SRM. A week later the rats that previously received saline now received CNO/OXTR antagonist and vice versa and were tested on the long-term social discrimination task to assess long-term SRM. In all experiments, the order of the discrimination tests (i.e. short-term and long-term social recognition memory), was randomized between cohorts. Since the same animals were being used as within subject controls (saline vs CNO or saline vs OXTR antagonist), a priori condition was used wherein animals that failed to exhibit short or long-term SRM on saline treatment were not considered for further analysis. This was determined based on a threshold of 0.05 on the ratio of duration of investigation (RDI) index calculated as (Investigation Time^Novel^ – Investigation Time^Familiar^) / (Investigation Time^Novel^ + Investigation Time^Familiar^).

### Viral Vectors

For specific silencing of OXT neurons, we used a previously validated AAV1/2-OXTp-hM4Dgi-mCherry virus, which has been shown to reduce mean frequency and input resistance of OXT neurons^32^. A control virus (AAV1/2-OXTp-mcherry) that lacks the DREADD backbone was used to account for the non-specific effects of CNO. To identify OXT neuron projection fibers from the PVH or SON, we used an anterograde virus driven by an OXT promoter (AAV1/2-OXTp-Venus)^33^. All OXTp viruses were produced and validated by Dr. Valery Grinevich’s laboratory at the Central Institute of Mental Health, University of Heidelberg, Germany. To identify SuM to CA2 projection neurons, a retrograde virus (AAV-Retrograde-Cre, Addgene #55636) and a cre-dependent virus (AAV9-DIO-Ef1a-eYFP, Addgene #27056) were used.

### Stereotaxic Surgery

Animals were anesthetized with 4% isoflurane for induction of anesthesia and maintained at 2% isoflurane and 2% oxygen using a tabletop vaporizer. The surgical area was shaved and aseptically cleaned. An incision was made along the dorsal midline of the skull, bregma and lambda were identified, the region of injection was marked, and a small burr hole (50um) was drilled. The virus (AAV1/2-OXTp-hM4Dgi-mCherry or AAV1/2-OXTp-Venus or AAV1/2-OXTp-mCherry) was loaded into a 20μl NanoFil syringe fitted with a 33gauge needle (cat no. NF33BL, World Precision Instruments Inc, Sarasota, FL, USA). A final volume of 270nl was injected into the PVH (A-P −1.7mm, M-L ±0.3mm, D-V 8.0mm) at a 10° angle. For CA2 (A-P −3.5mm, M-L ±4.2mm, D-V 3.3mm) injections, a volume of 0.05ul was used and for SuM (A-P −4.5mm, M-L ±4.2mm, D-V 8.9mm, 15° angle) injections, a volume 0.07ul was used. Following injection, the syringe was left in place for 10 min and withdrawn at a rate of 0.2 mm/min. The incision wound was closed using wound clips (EZ Clips, Stoelting Inc, Wood Dale, IL, USA). Rats received intraoperative subcutaneous fluids for hydration (Lactated Ringer Solution, Thermo Fisher Scientific, Waltham, MA, USA) and buprenorphine (0.05mg/kg) for analgesia. Additional analgesia (buprenorphine, 0.05mg/kg) was administered subcutaneously every 12h for 72h post-operatively. For cannulation, a guide cannula (7mm, P1 Technologies Inc, Roanoke, VA, USA) was implanted at a 15° angle (A-P −4.65mm, M-L – 0mm). Two additional bone screws (Stoelting Inc, Wood Dale, IL, USA) were implanted on the skull surface for anchoring, and the guide cannula was secured using dental cement (Stoelting Inc, Wood Dale, IL, USA). A dummy cannula (7mm) was placed inside the guide cannula and was left in place until the day of experiment when OXTR antagonist/saline was delivered using an infusion cannula (9mm). Animals were allowed to recover for 3 days before experimentation.

### Drugs

For DREADD experiments, clozapine N oxide dissolved in 0.05% DMSO and 0.9% saline or 0.9% saline (+0.05% DMSO) was injected intraperitoneally (i.p) using a 1ml BD Luer-Lok syringe (cat. No. 309328, BD Biosciences, Mississauga, Ontario, CA). For OXTR antagonist experiments, a stock (1mg/ml) of OXTR antagonist (desGly-NH_2_-d(CH_2_)_5_[D-Tyr^2^,Thr^4^]OVT was prepared by diluting the drug in 0.9% saline. A working stock of 0.25ug/ul was prepared on the day of the experiment and was injected using a syringe pump (Amuza Inc, San Diego, CA, USA). A 5ul Hamilton syringe (Hamilton Company, Reno, NV, USA) was connected to a plastic tubing on one end and an infusion cannula on the other. Saline/OXTR antagonist (0.25ug/ul) was loaded into the infusion cannula and a volume of 0.3ul (75ng total) was injected at a rate of 0.1ul/min. The specific dose was chosen based on studies demonstrating its effectiveness in regulating fear memory in rats^34,35^. The infusion cannula was left in place for 1min before being withdrawn.

### Short and long-term discrimination task

Short and Long-term SRM were assessed using previously published paradigms, known as social discrimination tasks^14,36,37^. Briefly, the paradigms involve an initial encounter with a social stimulus followed by a short (30min) or long (24hour) inter-trial interval after which the test animal is simultaneously exposed to the same stimulus (“Familiar”) as before and a novel stimulus (“Novel”). Social recognition is considered to have occurred when the test rat shows greater preference for the novel stimulus over the previously encountered stimulus^38^. For all behaviors, test and stimulus rats were habituated to handling and to the testing arena for 4 days before testing. To assess short-term SRM, test rats were placed in the testing arena (50×50×40cm) at the start of the experiment. 30min later, they received an i.p injection of either saline or CNO (8mg/kg) and 30min following the injection, a juvenile rat of a different strain (3-5 week old, Wistar or Wistar Hannover strain) was placed in an enclosure and introduced into the testing arena for the test rat to investigate for 5min (1^st^ encounter). Following an inter-trial interval of 30min, during which time the test rat remained in the testing arena, the juvenile rat from the 1^st^ encounter (“Familiar”) and a new juvenile rat (“Novel”) were placed in two small enclosures and introduced in two opposing corners of the testing arena for the test animal to investigate for 10min (“2^nd^ encounter”). The strain of the juvenile rats used for the first encounter were randomized such that each test rat interacted with a different strain across treatment sessions. Additionally, the position of the juvenile rat within the testing arena during the first encounter was always different than the one during the second encounter. To assess long-term-SRM, test rats received an i.p injection of either saline or CNO (8mg/kg) and were placed in the testing arena. 15min later, a juvenile rat of a different strain (3-5 week old, Wistar or Wistar Hannover strain) was placed in the testing arena for the test rat to freely interact and investigate for 1h (1^st^ encounter). After a 24h inter trial interval, the test rat was placed in the testing arena for 1h for habituation followed by introduction of the “Familiar” and a “Novel” rat that were placed in two small enclosures and introduced in two opposing corners of the testing arena for the test animal to investigate for 10min (2^nd^ encounter). Stimuli rats of different strains were used to enhance SRM acquisition, especially for long-term SRM^37^. In order to minimize the influence of spatial memory, the positioning of the stimuli rats during the 2^nd^ encounter was always different from the 1^st^ encounter (short-term SRM) and randomized between test subjects in both the short and long-term SRM paradigm.

### Novel Object Recognition Memory

Novel object recognition task was performed based on a previously established protocol^39^. Test rats were injected with 0.9% saline or CNO (8mg/kg). 15 min later, test rats were allowed to interact for 3 min with two identical objects (Lego or Cone), which were placed on one side of the arena (“1^st^ encounter”). After an inter-trial interval of 30min during which the test rat remained in the testing arena, test rats were introduced to one of the objects from the first encounter (“Familiar”) and a novel object (“Novel”) and was allowed to interact with both for 3min. The choice of objects was randomized across treatment groups.

### Behavioral Analysis

All behaviors were scored and quantified using TrackRodent, an open-source Matlab based automated tracking system that uses a body-based algorithm^40,41^. The traces and heat-maps were also obtained using the same system. The source code can be accessed on GitHub (https://github.com/shainetser/TrackRodent).

### Histology

Rats were anesthetized with an i.p injection of Ketamine (100mg/kg) and Xylazine (13mg/kg). Once a surgical plane of anesthesia was achieved, rats were peristaltically perfused at a rate of 30ml/min with 0.2M Sodium phosphate buffer for 6 min followed by 4% paraformaldehyde (PFA) at 40ml/min for 6 min. Brains were removed, immersed in 4% PFA overnight at 4°C, then placed in 30% sucrose in 1xPBS for 48h. Brains were flash frozen in a slurry of dry ice and isopentane and sectioned on a cryostat (Leica CM 1860 Leica Biosystems, Buffalo Grove, IL, USA).

### Immunohistochemistry

To visualize overlap between the DREADD virus and OXT neurons a total of 12 sections spanning the entire PVH were used. Briefly, sections were washed (3×10 min each in 1xPBS, 0.05% Triton X-100), blocked and permeabilized for 1h in 5% donkey serum in 1XPBS-0.5% triton-X-100 for 1h at room temperature (RT). They were then co-stained with anti-OXT (1:1000) and anti-DsRed (1:1000) antibodies for 24h at 4°C in 1xPBS-0.5% triton-X-100. Sections were then washed and incubated in donkey anti-mouse IgG 488 (1:1000) and donkey anti-rabbit IgG 594 (1:1000) in 0.5% Triton X-100 in PBS for 2h at RT. Sections were washed and mounted with antifade mounting Medium with DAPI.

For immunofluorescence experiments, PVH and SuM sections from an 8 week male SD rat were blocked in 5% donkey serum at room temperature (RT) and co-stained with OXT (1:1000) and calretinin (1:2000) or parvalbumin (1:2000) for 24h at 4°C. Sections were then washed and incubated in donkey anti-mouse IgG 488 (1:1000) and donkey anti-rabbit 594 (1:1000) or donkey-anti-goat 594 in for 2h at RT. Similarly, PVH and SuM sections from an OXTp-Venus injected rat were co-stained with anti-GFP (1:1000) and anti-calretinin (1:2000) or anti-parvalbumin (1:2000) antibodies and incubated for 24h at 4°C. This was followed by incubation in donkey anti-chicken IgG 488 and donkey anti-rabbit 594 or donkey anti-goat IgG 594 for 2h at RT. To demonstrate that the retrograde virus targeted the hippocampal CA2 region, hippocampal sections were incubated in anti-Cre (1:2000) and PCP-4 (1:2000). To identify CA2 projecting SuM neurons, SuM sections were incubated with anti-GFP (1:2000), anti-cre (1:2000), and anti-calretinin (1:2000) and were processed using similar incubations times as listed above.

To visualize OXT fibers using enzymatic staining, brain sections (40um) representing the PVH, SON or SuM from an 8 week male SD rat were used. Sections were treated with 3% hydrogen peroxide (H_2_O_2_) followed by 1h incubation in 5% goat serum. Next, sections were incubated with anti-OXT antibody (1:1000) for 40h at 4°C, washed and incubated with goat anti mouse HRP 2h at RT, and developed using an ImmPACT diaminobenzidine (DAB) peroxidase substrate. Alternate SuM sections from the same animal were used for staining tissue with 1% cresyl violet. Similarly, to visualize OXTp-Venus fibers, OXTp-Venus was injected into the PVH or SON. PVH, SON or SuM sections were treated with 3% H_2_O_2_ to block endogenous peroxidase activity and incubated in 5% goat serum followed by incubation with anti-GFP antibody for 40h at 4°C. Sections were washed and incubated in the goat anti chicken HRP for 2h at RT and developed using DAB. To examine infusion cannula localization for OXTR antagonist experiments, tissue sections were treated with 1% cresyl violet solution.

### RNAscope

Rat *Oxtr*^42^, *vglut2 (slc17a6)*^43^ and *vgat1 (slc32a1)*^43^ probes were purchased from ACDBio. Fresh brains were collected by cervical decapitation and flash frozen in a slurry of isopentane and dry ice. Tissue was immediately sectioned at 15um, mounted on glass slides (SuperFrost Plus Microscope Slides, Fisher Scientific) and frozen at −80°C until the day of experiment. RNAscope was performed using the following the manufacturer’s protocol (RNAscope Multiplex Fluorescent Reagent Kit, ACDBio, Newark, CA). Briefly, tissue sections were thawed at RT for 10min, fixed with 4% PFA for 15min at 4°C, and then dehydrated with ethanol at RT. Sections were then incubated in H_2_O_2_ for 10min and a mix of *Oxtr, vglut*2 and *vgat1* probes was added to the sections and left to incubate for 2h in a 40°C oven (HybEZ II Hybridization System, ACDBio, Newark, CA). This was followed by an amplification step that involved amplification probes (Amp1, Amp2, and Amp3), provided with the kit, and then an incubation step with opal dyes (Akoya Biosciences, Marlborough, MA) 520, 570, and 690 to visualize the RNA transcripts.

### Microscopy and Image Analysis

PVH sections (10-12) from OXT-hM4Dgi-mcherry injected rats were imaged on a confocal microscope (Leica SP5 DMI, Leica Micro-Systems, Buffalo Grove, IL, USA) at the Microscopy and Advanced Bioimaging CoRE at the Icahn School of Medicine at Mount Sinai. Sections were imaged at 20x and Z stacks were acquired at step size of 1.0um and stacked images were exported to FIJI (ImageJ) and single plane images were generated using Z project (maximum intensity projection)^44^. mCherry overlap with OXT neurons was manually counted from 3 animals (12 PVH sections each) and presented as % mCherry^+^/OXT^+^ (**Supplemental Fig. 1a**). Fluorescent in situ hybridization (RNAscope) images were acquired on a Zeiss AxioImager Z2M with ApoTome.2 at 10x, 40x and 63x magnification. Images were imported into FIJI and a grid drawn over the acquired image. Individual neurons were counted grid by grid using the cell counter plugin on FIJI (n=3 rats, 1 section/rat). DAB stained sections were acquired using a bright field microscope (EVOS, Thermo Fisher Scientific, Waltham, MA).

### Statistical analysis

Statistical analysis was performed using GraphPad prism 9.0 software (GraphPad Prism, San Diego, CA). Total investigation time between Familiar and Novel social stimuli were evaluated using a two-way repeated measures Analysis of Variance (RM-ANOVA) to compare main effects of treatment (Saline vs. CNO or Saline vs. OXTR antagonist) and social preference (Familiar vs. Novel). Sidak’s multiple comparison test was used for post-hoc testing.

### Reagents

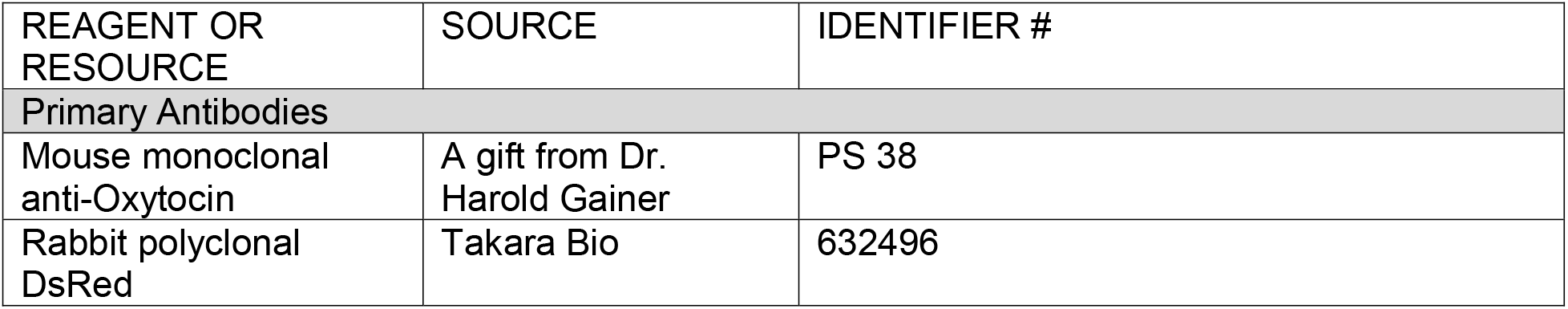
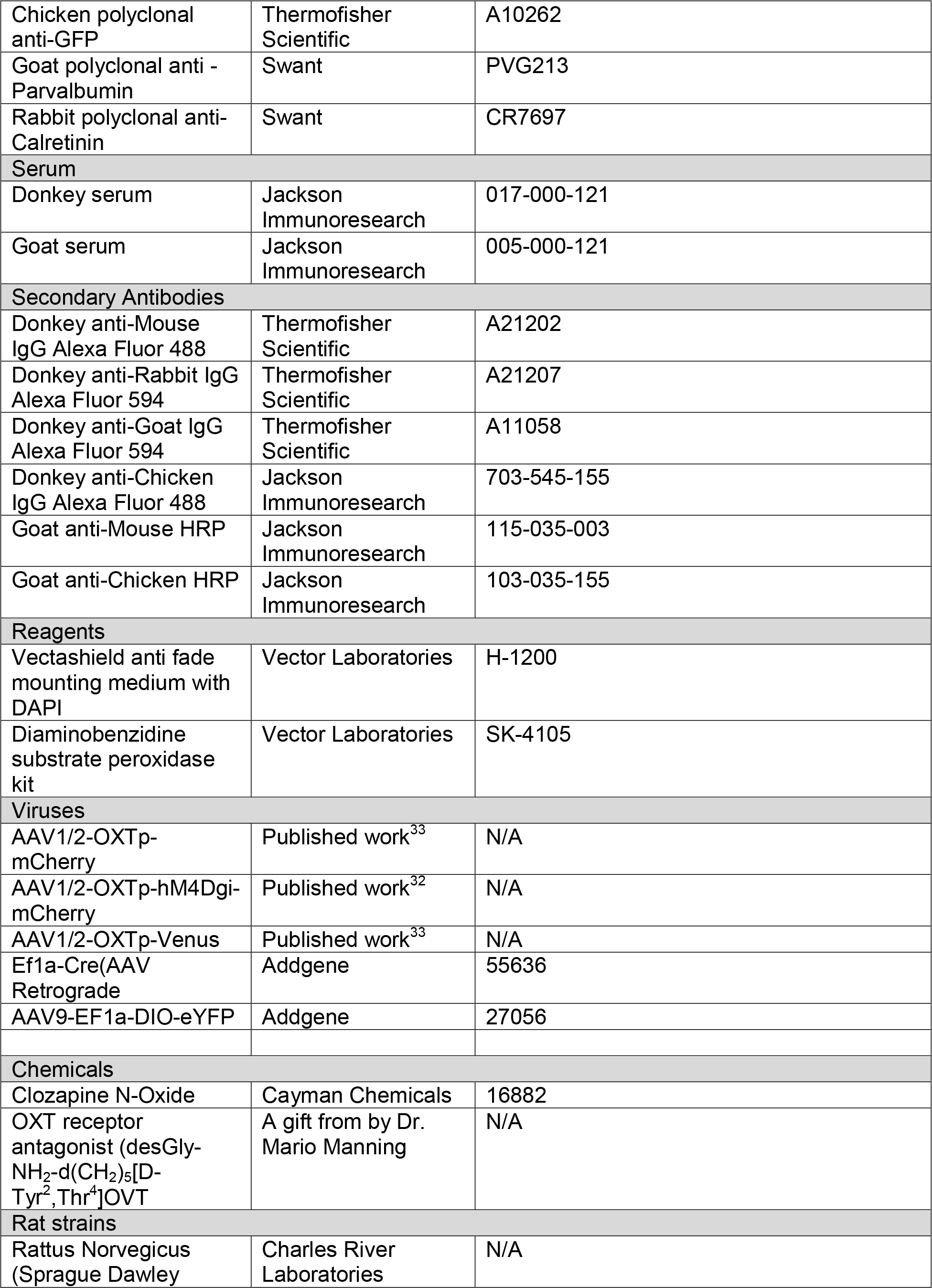
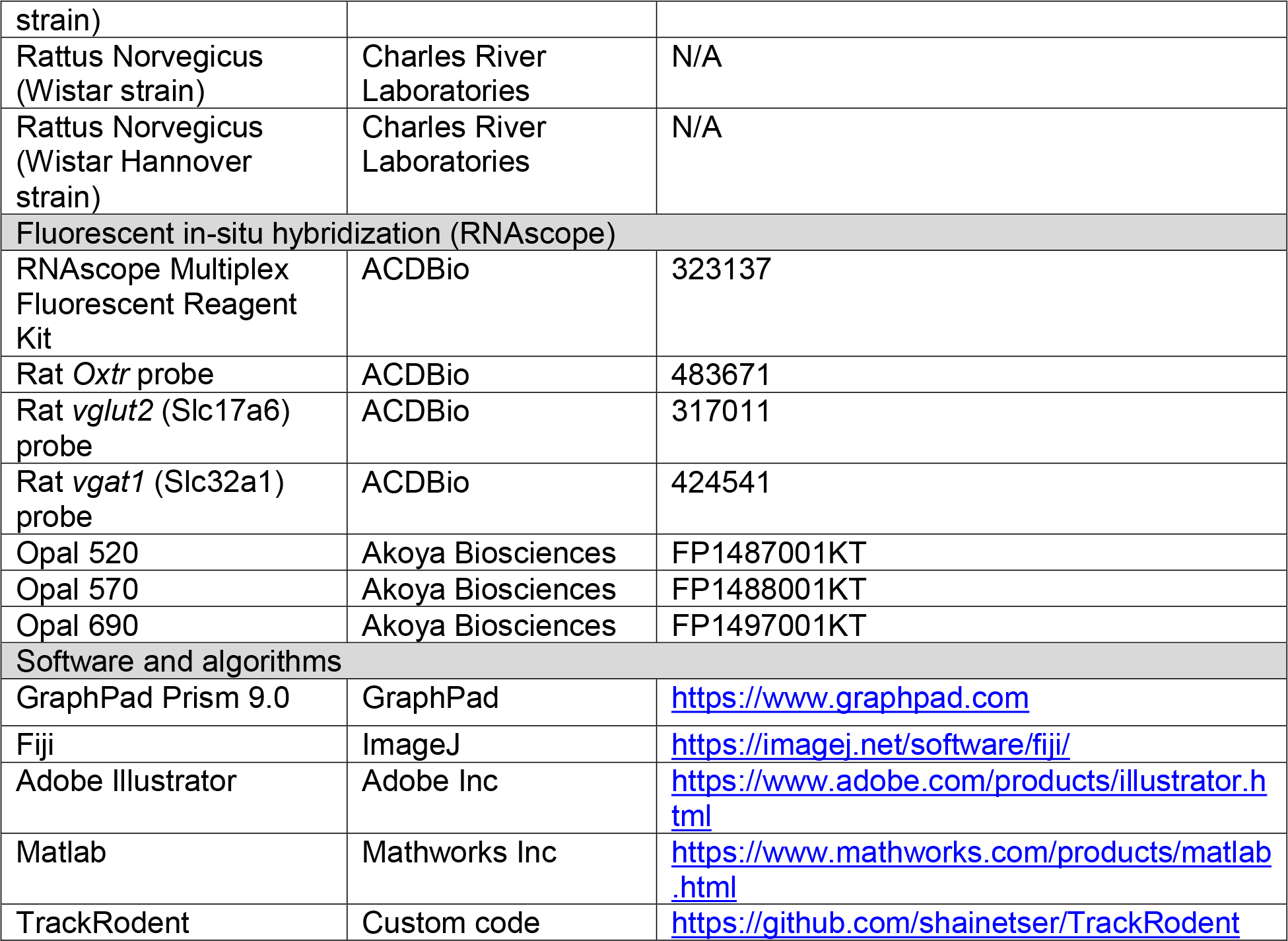

## ACKNOWLEDGEMENTS

This work was supported by the Seaver Foundation for Autism Research and Treatment (H.H.N, K.T.R, M.B), the National Institute of Mental Health (R01MH116108, H.H.N), a trainee pilot grant from the Mindich Child Health and Development Institute (K.T.R), Young Investigator Award from the Brain and Behavior Research Foundation (K.T.R) and the National Institute of Mental Health Diversity Supplement Award (MH116108-03S1, H.H.N, N.C). A.L. is a recipient of Marie Curie-Skladowska fellowship. V.G was supported by the German Research Foundation (DFG) grants GR 3619/15-1, GR 3619/16-1, SFB Consortium 1158-2 (V.G.), and GR 3619/13-1 and 3619/19-1 (S.W. and V.G.). S.W was supported by ISF-NSFC joint research program (grant No. 3459/20), the Israel Science Foundation (ISF grant 1361/17), the Ministry of Science, Technology and Space of Israel (Grant No. 3-12068) and the United States-Israel Binational Science Foundation (BSF grant No. 2019186, S.W and H.H.N). The authors would like to acknowledge Ms. Amanda Leithead and Ms. Michelle Kim’s support in reviewing the manuscript and thank the Microscopy and Advanced Bioimaging CoRE core at the Icahn School of Medicine for their guidance on imaging. Parts of Figs **1** and **6** were created with https://BioRender.com.

## AUTHOR CONTRIBUTIONS

K.T.R and H.H.N conceptualized and designed the study. K.T.R performed all DREADDS and OXTR antagonist experiments, brain surgeries, viral injections, RNAscope, and acquired all imaging data. M.B performed brain surgeries and viral injection for the OXT projection experiments. K.N helped on the behavioral analysis. N.C helped with the quantification of immunohistochemical data. A.F and V.G cloned and packaged the OXT-specific viruses. S.N and SW provided support on the open source behavioral analysis software. K.T.R and H.H.N, interpreted the data, prepared figures and wrote the manuscript. H.H.N, S.W, S.N and V.G helped interpret the data and revise the manuscript. All authors read and approved the final manuscript.

