## Supplementary material for "Oxytocin Activity in the Paraventricular and Supramammillary Nuclei of the Hypothalamus is Essential for Social Recognition Memory in Rats": Supplmental data

**Title**

**SUPPLEMENTAL INFORMATION**

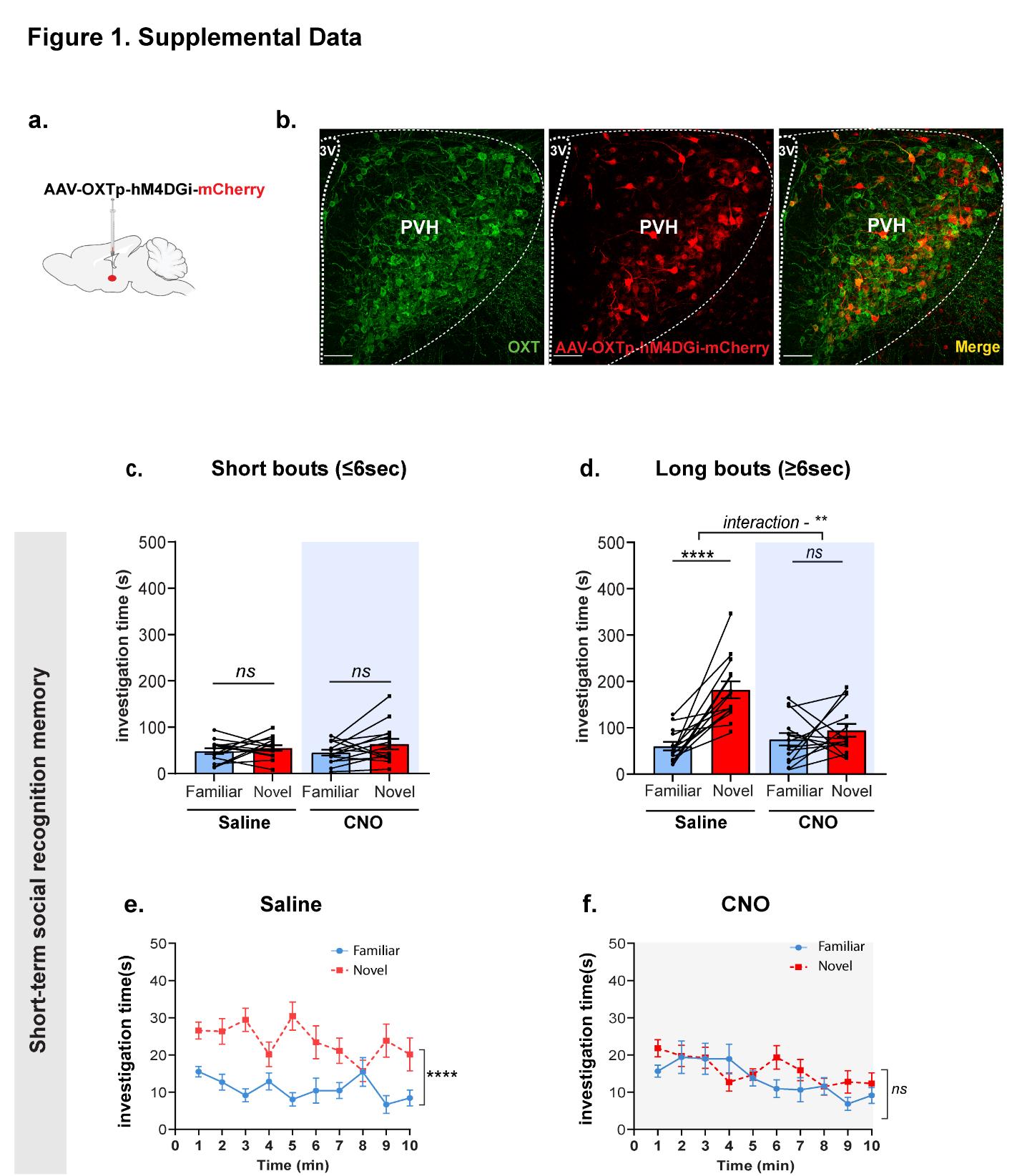

**Supplemental data Figure 1. Chemogenetic silencing of PVH-OXT neurons impairs short-SRM. a.** A schematic of the viral injection **b.** A representative image of PVH-OXT neurons showing AAV1/2-OXT-hM4DGi-mcherry and OXT co-expression in the PVH. **c.** Total investigation time of the novel vs. familiar stimuli during the 2^nd^ encounter for short bouts of interaction (≤6sec) during short-term SRM. No significant differences in the preference for novel over familiar stimuli following Saline or CNO injection (two-way, RM ANOVA, effect of treatment (Saline vs. CNO) x social preference (Familiar vs. Novel) interaction (F_1,26_ = 0.330, *P*=0.570, *ns*))*,* effect of treatment (F_1,26_ = 0.418, *P*=0.523, *ns*), and effect of social preference (F_1,26_ = 2.074, *P*=0.161, *ns*). **d.** Total investigation time of the novel vs. familiar stimuli during the 2^nd^ encounter for long bouts of interaction (≥6sec). There was a significant difference in the preference for novel over familiar stimuli following saline, whereas no difference was observed in preference for novel or familiar stimuli after CNO (treatment x social preference interaction (F_1,26_ = 10.51, ***P*=0.0032)), effect of treatment (F_1,26_ = 5.5, **P*=0.026), and effect of social preference, (F_1,26_ = 26.75, *****P*<0.0001). Post-hoc Sidak multiple comparison test, Saline (Familiar vs. Novel, ^****^*P*<0.0001) and CNO (Fam vs. Nov, *P=*0.424*, ns*)*.* **e.** Investigation time of novel vs familiar stimuli across time following Saline or CNO during short-term SRM. Saline group show consistent preference for novel over familiar stimuli across time (two-way RM ANOVA, time x social preference interaction (F_9,234_ = 2.36, **P=*0.01), effect of social preference (F_1,26_ = 44.71, *****P<0.0001*), and effect of time (F_9,234_ = 1.2, *P=*0.29). **f.** There was no clear preference for familiar or novel stimuli across time following CNO (two-way RM ANOVA, time x social preference interaction (F_9,234_ = 1.500, *P*=0.148, *ns*)), effect of social preference (F_1,26_ = 1.178, *P=*0.28, *ns*), and effect of time (F_9,234_ = 4.135, ***P<0.0001). SRM, Social recognition memory, PVH, paraventricular hypothalamus, OXT, oxytocin, CNO, clozapine-N-oxide. 3V, 3^rd^ ventricle. Data represented as mean ± SEM. Scale bar (100um). Data represented as Mean ± SEM.

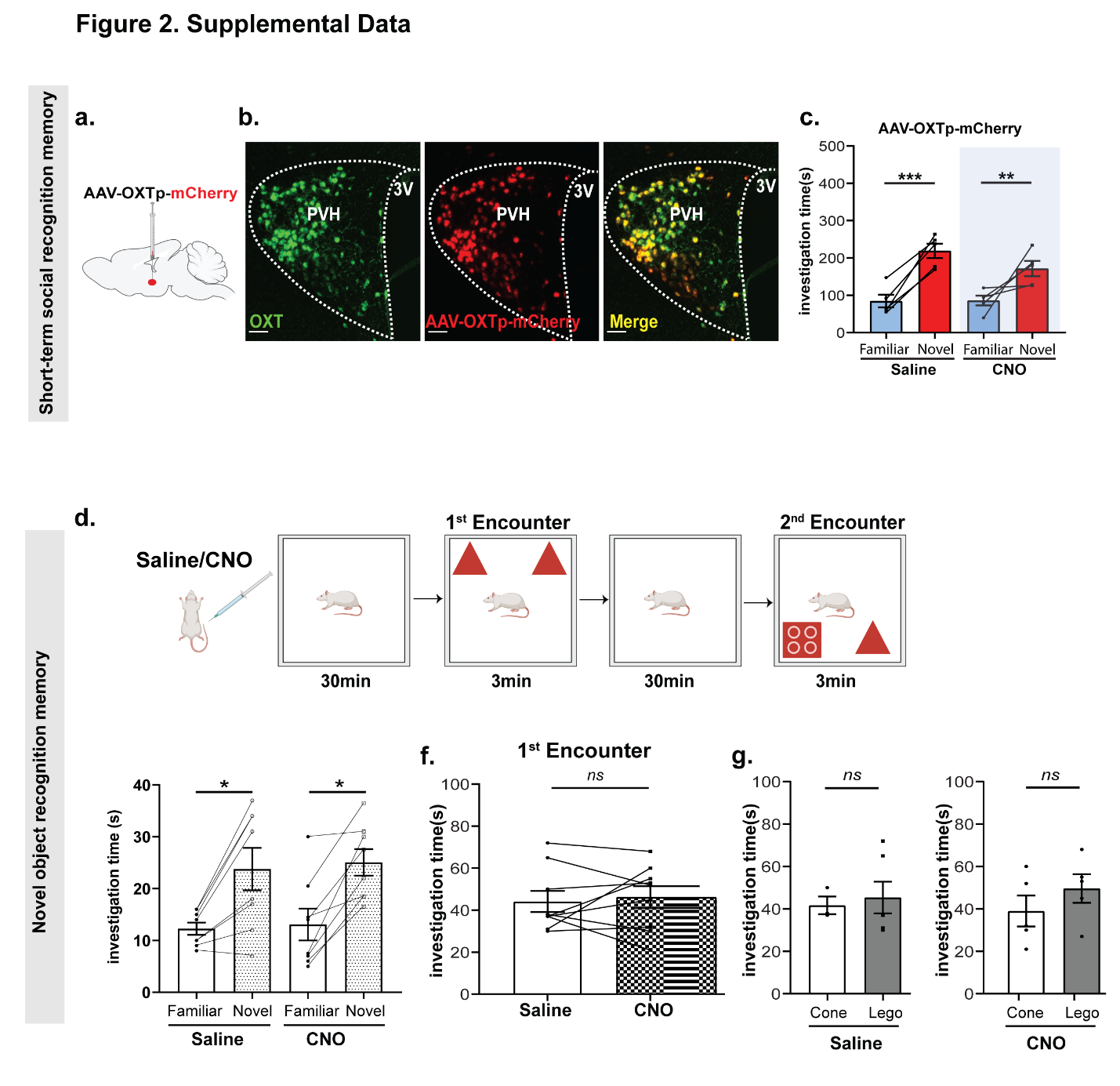

**Supplemental data Figure 2. CNO has no impact of SRM and chemogenetic silencing of PVH-OXT neurons does not impair object recognition memory. a.** A representative image showing overlap between AAV1/2-OXTp-mCherry (control) and OXT neurons in the PVH. **b.** Total investigation time of the novel vs. familiar stimuli during the 2^nd^ encounter in rats injected with the AAV1/2-OXTp-mCherry during S-SRM. Both saline and CNO treated rats showed a significant preference for the novel over the familiar social stimuli (two-way repeated measures (RM) ANOVA, social preference (Familiar vs. Novel) x treatment (Saline vs. CNO) interaction (F_1,8_ = 1.311, *P*=0.28, n=5), effect of social preference (F_1,8_ = 75.01, ***P*<0.0001), and effect of treatment (F_1,8_ = 1.15, *P*=0.31). Post-hoc, Sidak multiple comparison test, Saline (Familiar vs. Novel, ****P*=0.0001) and CNO (Familiar vs. Novel, ***P=*0.006). **d.** A Schematic of the novel object recognition paradigm. **e.** Total investigation time of the novel vs. familiar stimuli during the 2^nd^ encounter**.** Saline and CNO injected rats showed significant preference for the novel vs. familiar object (two-way RM ANOVA**,** treatment x social preference (Familiar vs. Novel) interaction, (F_1,14_ = 0.01, *P=*0.91, n=8)), effect of object preference (F_1,14_ = 11.27, ***P<*0.004) and effect of treatment (F_,1,14_ = 0.23, *P=*0.63). Post-hoc Sidak multiple comparison test revealed a significant difference in investigation time between familiar and novel object in both saline and CNO treated conditions saline (Familiar vs. Novel object), **P*=0.01) and CNO (Familiar vs. Novel object, **P=*0.014). **f.** Investigation time of the object stimuli during the 1^st^ encounter. There was no significant difference in the investigation time during the 1^st^ encounter between Saline and CNO injected groups (two-tailed paired student’s *t-*test, t_7_=0.52, *P*=0.61, *ns*). **g.** There was no innate preference for either of the two kinds of objects (Cone vs. Lego) following Saline (left graph) or CNO (right graph) injection (two-tailed unpaired t-test, Cone vs. Lego, Saline, t_7_=0.32, *P*=0.75, *ns,* Cone vs. Lego, CNO, t_8_ = 1.07, *P*=0.31, ns). S-SRM, Short-term Social recognition memory, PVH, paraventricular hypothalamus, OXT, oxytocin, CNO, clozapine-N-oxide. 3V, 3^rd^ ventricle. Data represented as mean ± SEM. Scale bar (100um). Data represented as Mean ± SEM.

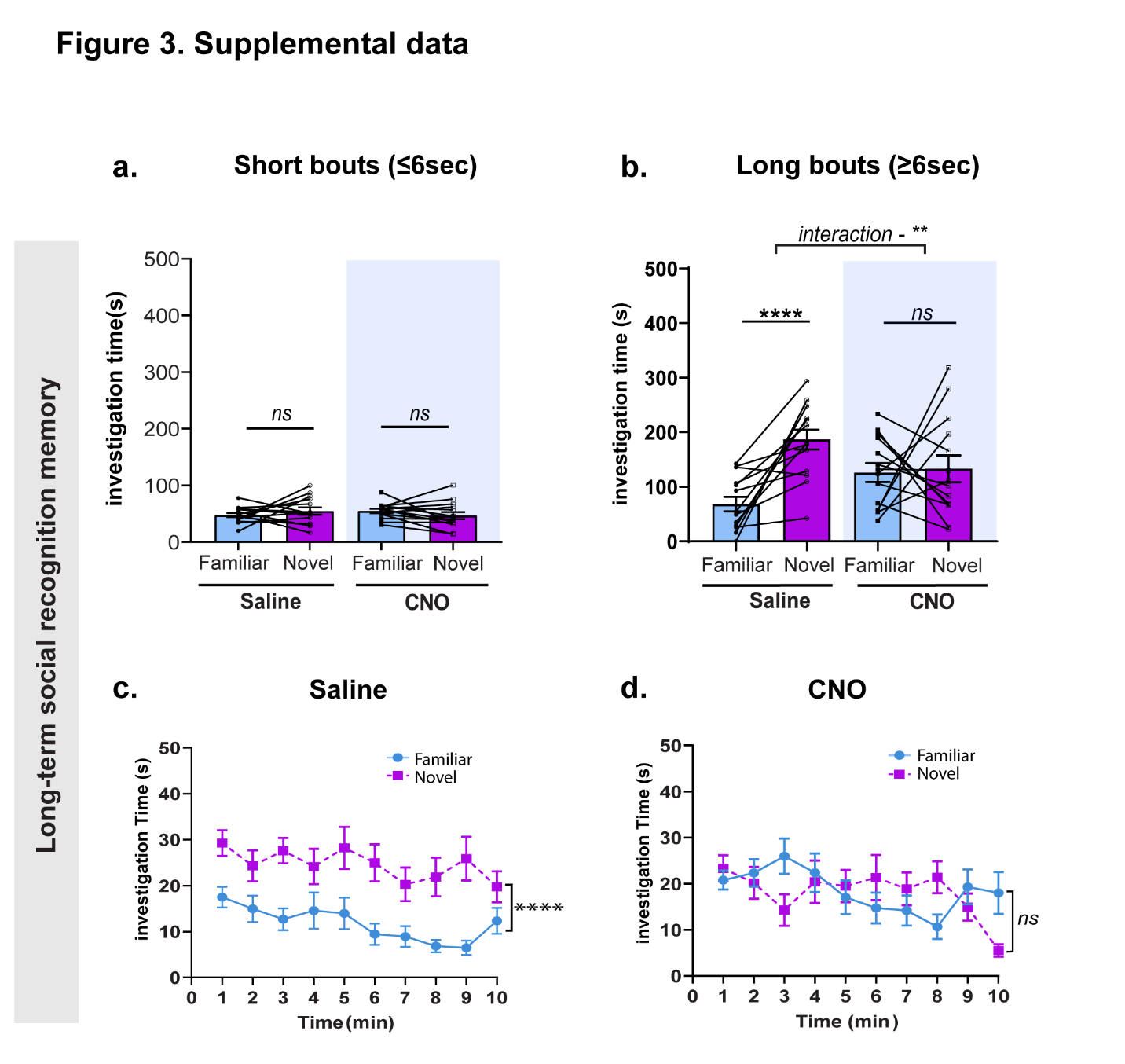

**Supplemental data Figure 3. Chemogenetic silencing of PVH-OXT neurons impairs long-term SRM.** Total investigation time of the novel vs. familiar stimuli during the 2^nd^ encounter for short bouts of interaction (≤6secs) during long-term SRM. No significant differences in the preference for novel over familiar stimuli following Saline or CNO (effect of treatment, F_1, 26_ = 0.002, *P*=0.957), effect of social preference (Familiar vs. Novel, F_1, 26_ = 0.01, *P*=0.920, treatment x social preference interaction, F_1, 26_ = 2.396, *P*=0.133). **b.** Total investigation time of the novel vs. familiar stimuli during the 2^nd^ encounter for long bouts of interaction (≥6secs). There was a significant difference in the preference for novel over familiar stimuli following saline, whereas no significant difference in the preference either stimuli was observed following CNO (treatment x social preference interaction, F_1,26_ = 8.08, ***P*=0.008) effect of treatment, (F_1,26_ = 0.011, *P*=0.913), effect of social preference, F_1,26_ = 12.19, ***P*=0.0017). Post-hoc Sidak multiple comparison test, Saline (Familiar vs. Novel, *****P*<0.0001) and CNO (Familiar vs. Novel, *P=*0.957*, ns*)*.* **c.** Investigation time of the novel or familiar stimuli across time following saline or CNO during long-term SRM**.** Saline injected rats showed a clear preference for the novel over the familiar stimuli across time, (two-way RM ANOVA, time x social preference interaction (F_9, 234_ = 0.7475, *P*=0.665, *ns*), effect of time, (F_9, 234_ = 2.038, **P*=0.036), effect of social preference (F_1, 26_ = 28.94, *****P*<0.0001). **d.** The same animals showed no clear preference for the either stimuli across time following CNO injection, (time x social preference (F_9, 234_ = 3.240, ****P=*0.001), effect of social preference (F_1, 26_ = 0.03546, P=0.852, *ns*), effect of time (F_9, 234_ = 2.236, **P*=0.020). L-SRM, Long-term Social recognition memory, Data represented as mean ± SEM.

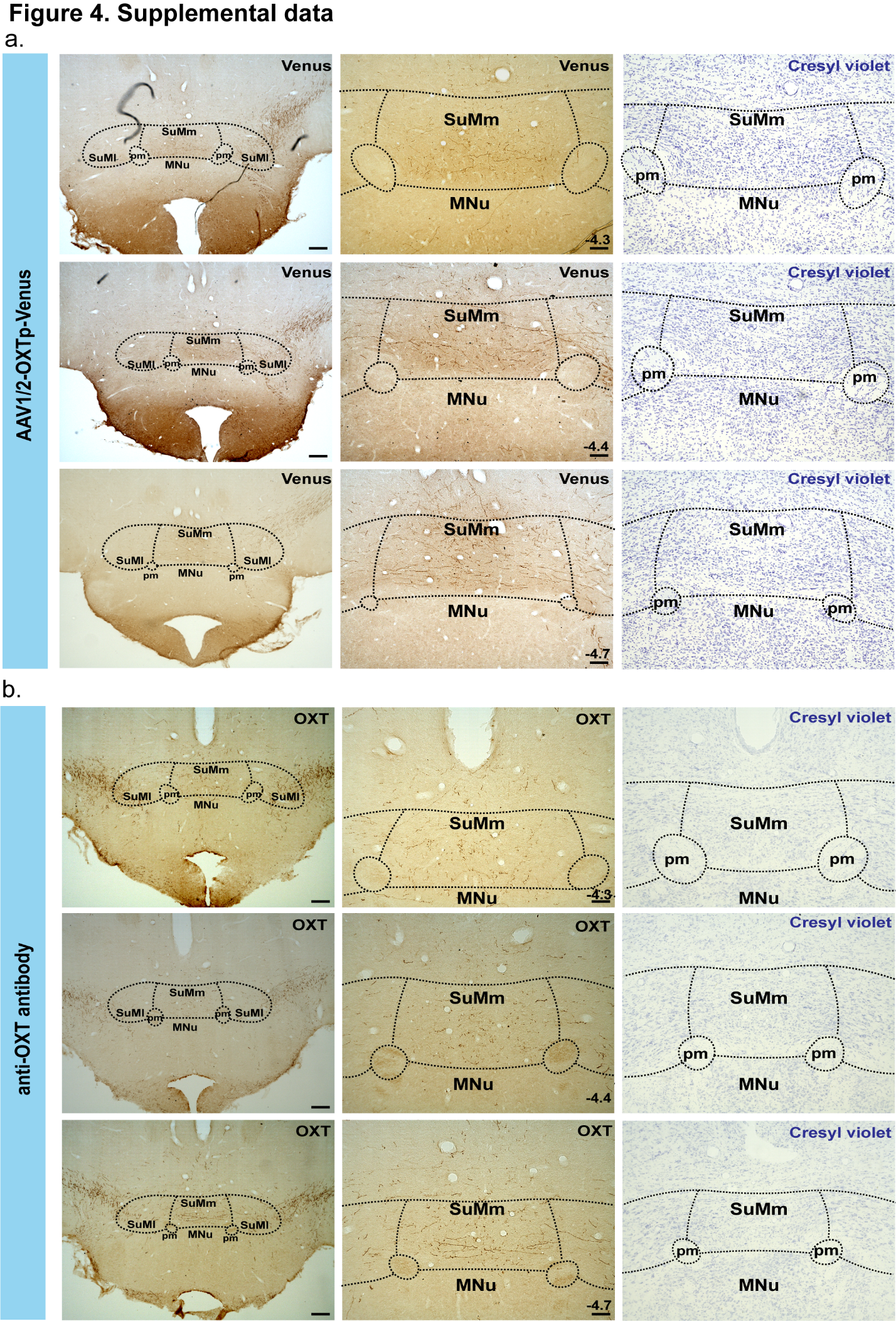

**Supplemental data Figure 4. OXT and Venus positive fibers are distributed through rostral-caudal aspects of the medial and lateral parts of the SuM. a.** Enzymatic labeling for Venus in rats injected with the AAV1/2-OXTp-Venus within the PVH, shows the distribution of Venus-positive fibers in the rostro-caudal aspect of the medial and lateral parts of SuM (Bregma -4.3 to -4.7mm). **b.** Enzymatic labeling for OXT in the SuM of wild type rats shows the distribution of OXT fibers in the rostro-caudal aspect of the medial and lateral parts of the SuM (Bregma -4.3 to -4.7mm). **Left and middle panels** are low (4x) and high magnification (10x) of the SuM, respectively, showing Venus-positive fibers (a) or OXT-positive fibers (b). Right panels are SuM sections stained with cresyl violet to highlight the SuM anatomy**.** Low magnification, scale bar 250um, high magnification, scale bar 100um. SuM, supramammillary nucleus, SuMl, lateral supramammillary nucleus, MNu, Mammillary nucleus, pm, principal mammillary tract. OXT, oxytocin.

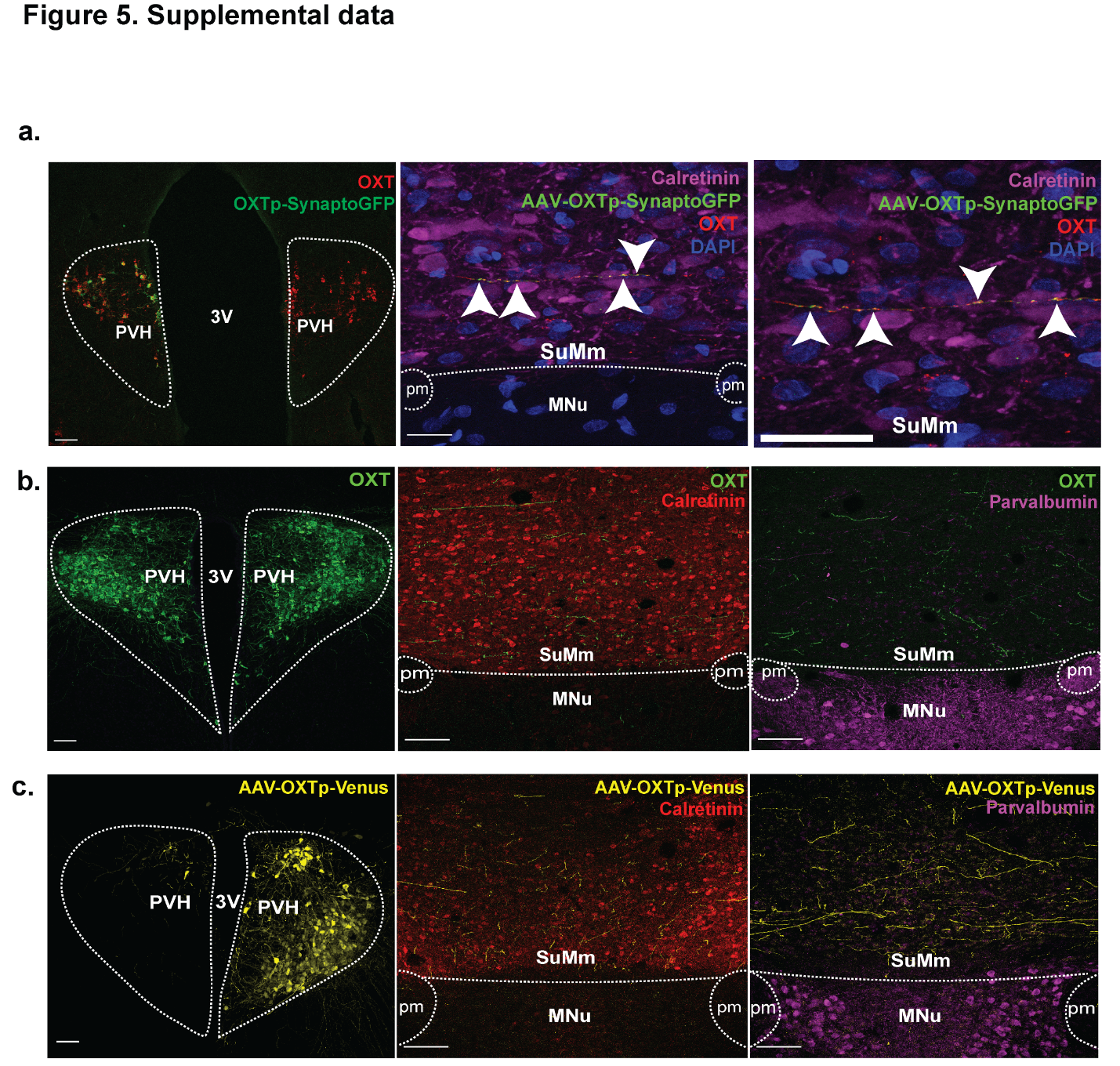
**Supplemental data Figure 5. PVH-OXT projection terminals are localized within the SuM. a. (Left)** A representative PVH section from a wild type rat, which was injected unilaterally with AAV1/2-OXTp-Synaptohpysin-GFP (green), co-stained with an anti-OXT antibody (red) to demonstrated the specificity of the virus**. (Middle)** lower (4x) and (**Right**) higher (10x) magnification of a SuM section showing presence of GPF positive puncta along the length of an OXT-labeled fiber **b.** Immunofluorescent labeling for OXT in the SuM of a wild type rat shows distribution of OXT fibers in the lateral and medial and the rostro-caudal (Bregma -4.3 to -4.7mm) aspect of SuM. (**Left**) A cross section of PVH stained with an anti-OXT antibody. (**Middle**) A cross section of the SuM co-labeled with an anti-OXT antibody to stain OXT fibers and an anti-calretinin antibody to highlight the SuM. (**Right**) A cross section of the SuM co-labeled with an anti-OXT antibody to stain OXT fibers and an anti-parvalbumin to distinguish SuM from the MNu. **c.** (**Left**) A representative PVH section from a wild type rat, which was injected unilaterally with AAV1/2-OXTp-Venus (yellow). (**Middle**) A cross section of the SuM showing the Venus distribution and co-labeled with an anti-calretinin antibody to highlight the SuM. (**Right**) A cross section of the SuM showing the Venus distribution and co-labeled with an anti-parvalbumin to distinguish SuM from the MNu. Low magnification, scale bar 250um, high magnification, SuMm, medial supramammillary nucleus, 3V, 3^rd^ Ventricle, MNu, Mammillary nucleus, pm, principal mammillary tract. OXT, oxytocin. Scale bar 100um (**a, b, c**, left, middle and right panels), 20um (**a**, middle panel) and 50um (**a**, right panel).

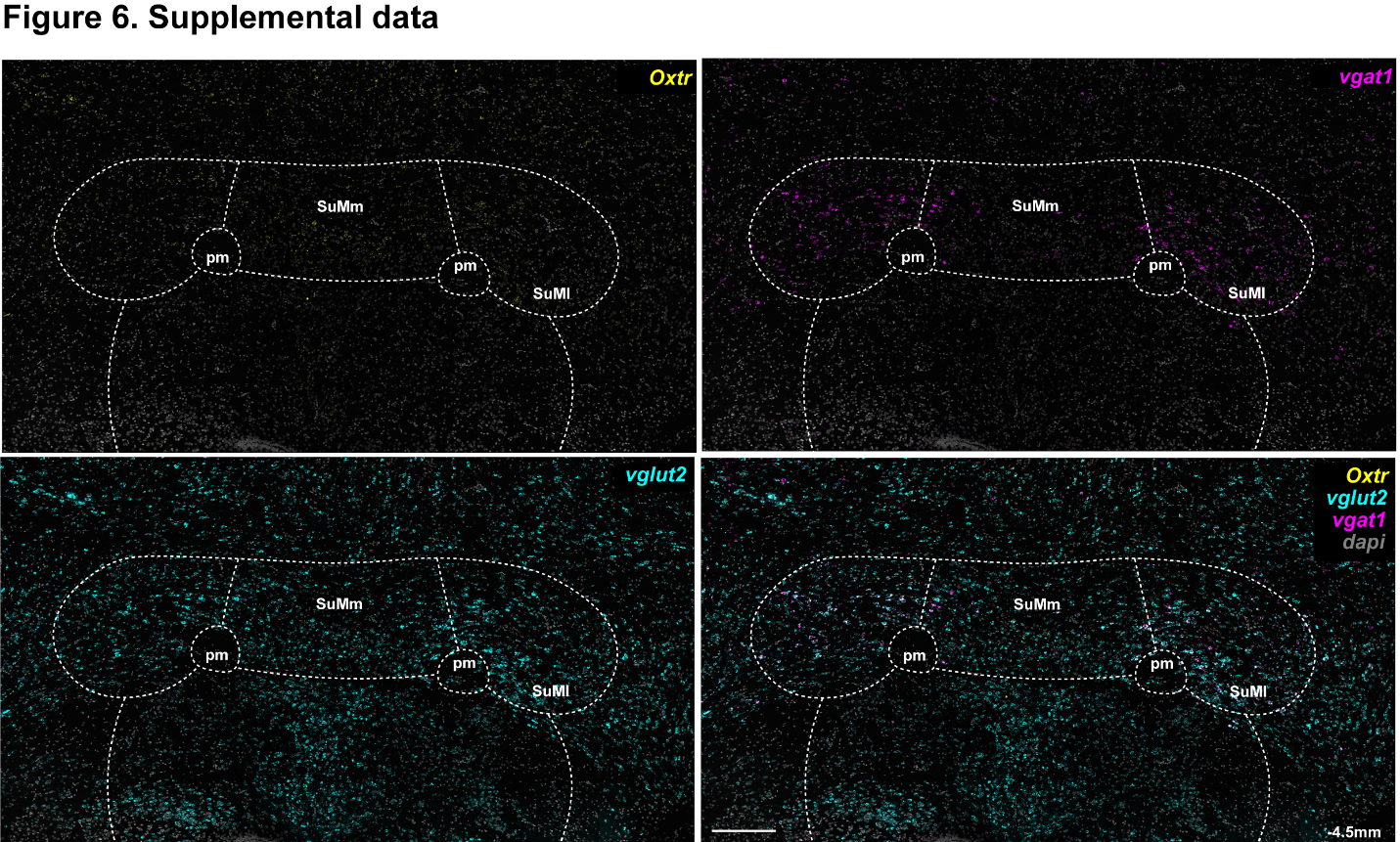

**Supplemental data Figure 6. In situ hybridization (RNAscope) using probes for *Oxtr*, *vglut2* and *vgat1* on a SuM section show OXTR distribution within the SuM.** A tiled image of SuM section following hybridization with *Oxtr*, *vgat1* and *vglut2* probes, show the expression pattern of *Oxtr*, *vglut2* and *vgat1* transcripts across the medio-lateral aspect of the SuM. Scale bar 1000um. SuMm, medial supramammillary nucleus, SuMl, lateral supramammillary nucleus MNu, Mammillary nucleus, pm, principal mammillary tract.

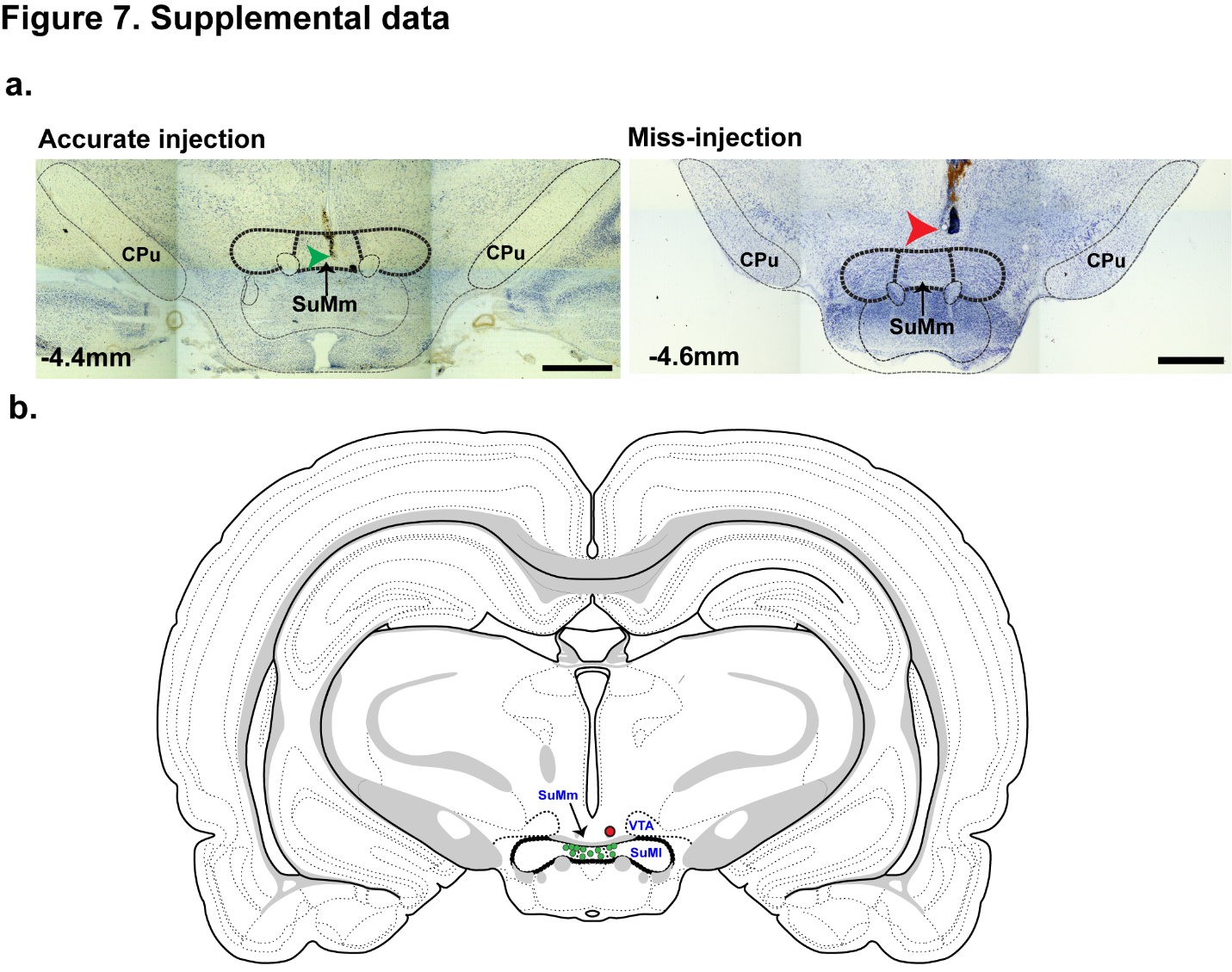
**Supplemental data Figure 7. Depiction of the drug cannula localizations in all rats that were tested with OXTR antagonist. a.** SuM sections from two independent rats implanted with cannulas to target the SuM are stained with cresyl violet to show an example of an accurate (left) and inaccurate (right) targeting of the SuM, as highlighted with the greed and red arrow, respectively. **b.** A modified Swanson rat atlas image (level 34) indicating correct (green circles) and incorrect (red circle) placement of the cannula tip within the SuM for each of the tested rats within the cohort. -4.4mm and -4.6mm denotes position of the section relative to bregma.

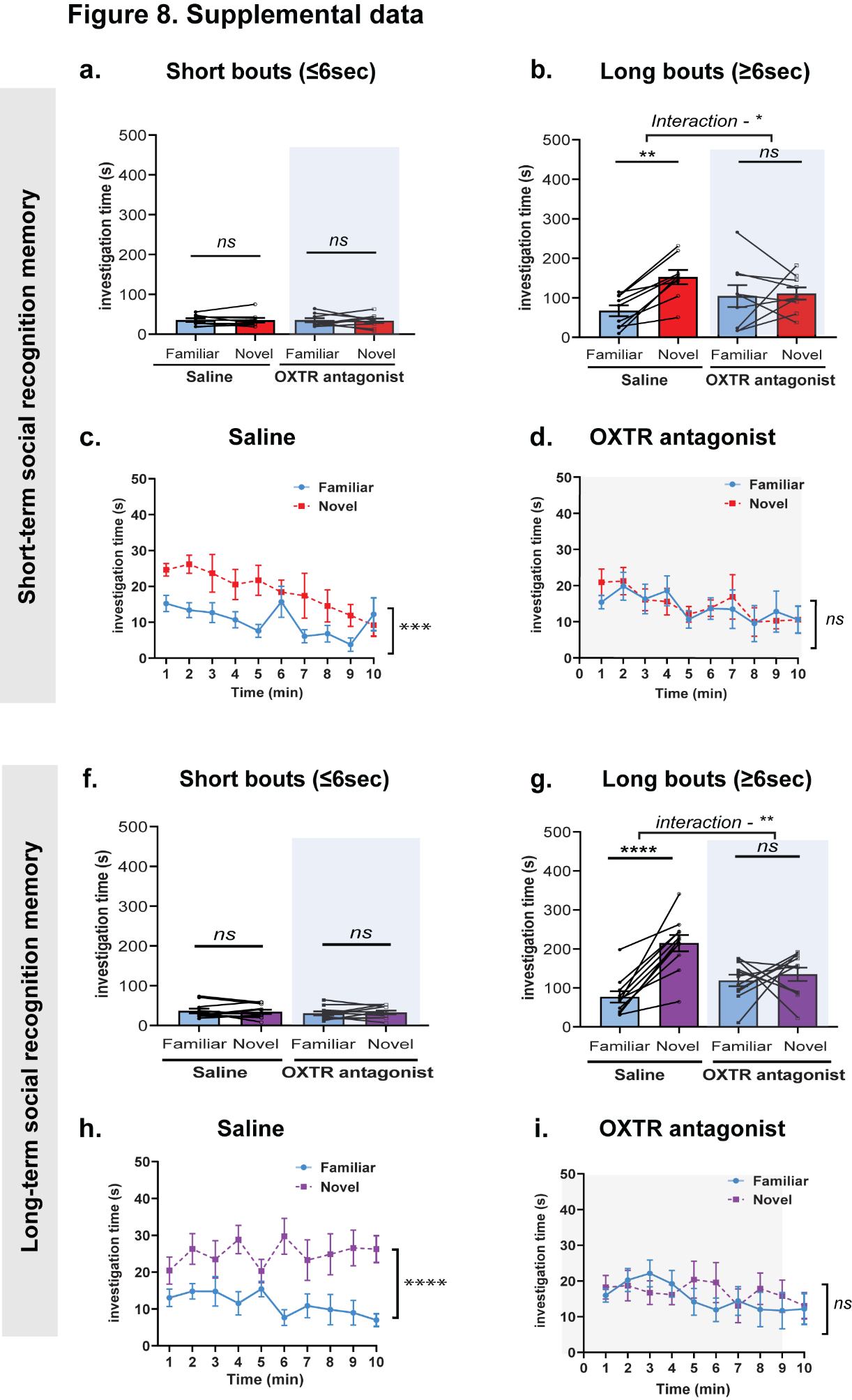

**Supplemental data Figure 8. OXTR antagonism in the SuM affects short and Long-term SRM. a.** Total investigation time for novel vs. familiar stimuli during the 2^nd^ encounter for short bouts of interaction (≤6sec) during short-term SRM. There was no significant difference for preference for novel over familiar following saline or OXTR antagonist infusion (treatment x social preference interaction (F_1,16_ = 0.03, *P*=0.86, *ns*), and effect of treatment (F_1,16_ = 0.04, *P*=0.83, *ns*), effect of social preference (Novel vs. Familiar, F_1,16_ = 0.019, *P=*0.89, *ns*) in short bouts. **b.** Total investigation time for the novel vs. familiar stimuli during the 2^nd^ encounter for long bouts (≥6sec). There was a significant difference in the preference for novel over familiar stimuli following saline, whereas the same animals showed no clear preference for novel or familiar stimuli following OXTR antagonist infusion (treatment x social preference interaction (F_1,16_ = 7.41, **P*=0.01), effect of treatment (F_1,18_ = 0.02, *P*=0.88), and effect of social preference (Novel vs. Familiar, F_1,16_ = 3.78, *P=*0.06). Post-hoc, Sidak multiple comparison test, saline (Familiar vs. Novel, ^**^*P*=0.008) and OXTR antagonist (Familiar vs. Novel, *P=*0.54*, ns*). **c.** Investigation time for familiar vs. novel stimuli across time following saline or OXTR antagonist infusion during short-term SRM. Saline infused group showed a clear preference for novel over familiar stimuli across time (two-way RM ANOVA, time x social preference interaction (Familiar vs. Novel) (F_9,144_ = 1.18, *P=*0.30*, ns*), effect of social preference, F_1,16_ = 16.85, ****P=*0.0009), and effect of time (F_5.3,84.83_=3.22, **P=*0.008). **d.** The same animals showed no clear preference for the familiar over the novel stimuli across time following OXTR antagonist infusion (two-way RM ANOVA, time x social preference interaction (F_9,144_ = 0.27, *P*=0.98, *ns*), effect of time (F_4.2,67.7_ = 2.2, *P*=0.07, *ns*), and effect of social preference (Novel vs. Familiar (F_1,16_ = 0.04, *P=*0.83, *ns*). **f.** Total investigation time of the novel vs. familiar stimuli during the 2^nd^ encounter for the short bouts of interaction (≤6sec) during long-term SRM. No significant differences for preference for novel over familiar following saline or OXTR antagonist infusion (treatment x social preference interaction (F_1,20_ = 0.60, *P*=0.44, *ns*), effect of treatment (F_1,20_ = 1.85, *P*=0.18, *ns*), and effect of social preference (Novel vs. Familiar, F_1,20_ = 0.0001, *P=*0.99, *ns*) in short bouts. **g.** Total investigation time of the novel vs. familiar stimuli during the 2^nd^ encounter for the long bouts of interaction (≥6sec). There was a significant preference for novel over familiar stimuli following saline infusion on long bouts, however the same animals after OXTR antagonist infusion did not show a clear preference for either stimuli (treatment x social preference interaction (F_1,20_ = 14.56, ***P*=0.0011), effect of treatment (F_1,20_ = 1.396, *P*=0.25, *ns*), and effect of social preference (F_1,20_ =18.42, ****P=*0.0004). Post-hoc, Sidak multiple comparison test, Saline (Familiar vs. Novel, ^****^*P*<0.0001) and OXTR antagonist (Familiar vs. Novel, *P=*0.76*, ns*). **h.** Investigation time for familiar or novel stimuli across time following saline or OXTR antagonist infusion during long-term SRM (two-way RM ANOVA, time x social preference interaction (F_9,180_ = 1.22, *P=*0.28, *ns*), effect of social preference (F_1,20_ =29.18, *****P<*0.0001), and effect of time (F_5.4,108.4_=0.30, *P=*0.91*, ns*). **i.** The same animals showed no clear preference for the familiar over the novel stimuli across time following OXTR antagonist infusion (two-way RM ANOVA, time x social preference (F_9,180_ = 0.62, *P*=0.77, *ns*), effect of time (F_6.20,124.0_ = 0.76, *P*=0.60, *ns*), and effect of social preference (Novel vs. Familiar, F_1,20_ = 0.4, *P=*0.53, *ns*).
